# A Transcriptomics-Based Computational Drug Repurposing Pipeline Identifies Simvastatin And Primaquine As Therapeutics For Endometriosis

**DOI:** 10.1101/2025.05.28.656743

**Authors:** Tomiko T. Oskotsky, Xinyu Tang, Erin Arthurs, Arpita Govil, Ferheen Abbasi, Arohee Bhoja, Daniel J. Bunis, Abby Lau, Jakob Einhaus, Maïgane Diop, Juan C. Irwin, Brice Gaudilliere, David K. Stevenson, Linda C. Giudice, Stacy L. McAllister, Marina Sirota

**Affiliations:** Bakar Computational Health Sciences Institute, University of California, San Francisco; San Francisco, CA, USA; Division of Clinical Informatics and Digital Transformation, Department of Medicine, University of California, San Francisco; San Francisco, CA, USA; Department of Gynecology and Obstetrics, Emory University; Atlanta, GA, USA; Center for Reproductive Sciences, Department of Obstetrics, Gynecology and Reproductive Sciences; University of California, San Francisco, CA, USA; Department of Anesthesiology, Perioperative and Pain Medicine, Stanford University; Stanford, CA, USA; Department of Pediatrics, Stanford University; Stanford, CA, USA; Department of Pediatrics, University of California, San Francisco; San Francisco, CA, USA

**Keywords:** bioinformatics, endometriosis, therapeutics, transcriptomics, drug repositioning

## Abstract

Endometriosis has limited treatment options, prompting the search for novel therapeutics. We previously used a transcriptomics-based computational drug repositioning pipeline to analyze public bulk transcriptomic data of eutopic endometrium from cases and controls and identified several drug candidates. Fenoprofen, our top in silico candidate, was validated in a rat model of endometriosis-associated pain. Building on this, we evaluated herein two additional candidates, simvastatin (a cholesterol-lowering drug) and primaquine (an antimalarial), based on strong endometrial gene expression reversal scores and favorable safety profiles. Using the rat model, we conducted behavioral testing, bulk RNA sequencing, and differential expression analysis to assess their therapeutic potential. We also assessed endometriosis diagnosis among patients prescribed simvastatin in electronic medical records (EMR) across six University of California (UC) healthcare institutions. In vivo validation using a rat model of endometriosis demonstrated that both simvastatin and primaquine significantly reduced vaginal hyperalgesia, a surrogate marker of endometriosis-related pain. RNA-seq of uteri and lesions confirmed reversal of disease-associated gene expression signatures following treatment. Analysis of UC-wide EMR data found lower relative risk of endometriosis among those prescribed simvastatin compared to a matched control group. Overall, simvastatin and primaquine attenuated pain-associated behaviors and reversed endometriosis-related gene expression changes in an animal model. Moreover, simvastatin prescription was associated with a lower relative risk of endometriosis in our retrospective multi-center cohort study. These findings highlight their potential as repurposed therapeutics for endometriosis and support the effectiveness of computational drug repositioning in identifying new treatment strategies.

**One Sentence Summary:** Simvastatin and primaquine reduced endometriosis pain and reversed gene signatures, with simvastatin also linked to lower disease risk.

## INTRODUCTION

Endometriosis is a common, life-altering disease, with symptoms including severe dysmenorrhea, chronic pelvic pain, and infertility(*1*). People with endometriosis can be undiagnosed for years, given non-specific symptoms that can overlap with other disorders and the requirement for surgical confirmation of disease(*1*). It has been suggested that 10% of reproductive-aged women have endometriosis(*2*), but this value may be an underestimate due to precise diagnostics requiring laparoscopy to identify and biopsy suspected endometriosis lesions. Approximately 12-32% of menstruating women who have surgery for pelvic pain and 50% who have infertility have been found to have endometriosis(*3*). The disease is not only a significant public health issue, but it also has a large economic impact on health expenditures of almost $70 billion annually in the U.S.(*4*).

Endometriosis is characterized by growth of endometrial tissue outside the uterus that responds to cyclic hormonal changes, similar to uterine endometrial tissue during a normal menstrual cycle(*2*). These lesions commonly exist at extrauterine sites such as the pelvic peritoneum, ovaries, and bowel, where they elicit an inflammatory response, fibrosis, and pain(*5*). The eutopic endometrium in women with endometriosis also differs from that of those without endometriosis, showing aberrant nerve fiber infiltration, increased angiogenesis, and disrupted progesterone response(*6, 7*). Although endometriosis was identified over 100 years ago, the disease remains with limited medical treatment options(*3*). Current evidence does not support any medication for primary prevention of endometriosis; however, as an estrogen-driven disease, hormonal therapies have been used to treat pain symptoms and reduce post-surgical recurrence(*2, 8–10*). First-line treatments for dysmenorrhea and non-menstrual pelvic pain associated with endometriosis typically include estrogen-progestin and progestin-only contraceptives or gonadotropin-releasing hormone (GnRH) analogues, as well as nonsteroidal anti-inflammatory drugs (NSAIDs)(*3, 8*). Unfortunately, existing medical therapies for endometriosis-related pain are often ineffective, with individuals experiencing minimal or transient pain relief or intolerable side effects limiting long-term use(*5*). Surgical excision of lesions can relieve symptoms, although combining lesion excision with hysterectomy has been shown to yield greater improvements in pain and quality of life(*11*). Even after surgery, many patients continue to experience symptoms(*12*). This underscores the need for new endometriosis treatment strategies that provide effective, non-hormonal control of pain, minimize local and systemic inflammation associated with pain and infertility, and, where possible, reduce the risk of developing disease or recurrence after surgery. Development of new drugs for endometriosis has been challenging due to the heterogeneity among affected individuals, variable symptoms, and complex etiologies, as well as historically lower investment in women’s reproductive health conditions(*13, 14*). Additionally, drug development for endometriosis has been limited as it is both costly and time-intensive, often requiring millions of dollars and decades of work, for therapeutics to finally become commercially available(*15*).

Computational drug repositioning methods allow for identification of new therapeutic applications for drugs already on the market in a fraction of the time and funds that it takes to develop and test entirely new drugs(*16*). Our group developed a method that identifies potential therapeutics that reversegene expression profiles of a disease -- i.e., genes downregulated in the disease are upregulated by the drug, and vice versa(*17*). This approach leverages transcriptomics data to generate genome-wide expression profiles from comparisons between disease and healthy samples or drug-treated and untreated cells(*17*). In the past, this method has been successfully applied to identify both known and novel treatments for inflammatory bowel disease(*18*), dermatomyositis(*19*), liver cancer(*20*), preterm birth(*21*), and Alzheimer’s disease(*22*). Our recent work to identify therapeutic candidates for endometriosis leveraged publicly available disease and drug gene-expression signatures in conjunction with a transcriptomics-based computational drug repositioning pipeline and validated the top candidate(*23*). Specifically, we applied this *in silico* approach to endometriosis disease signatures that were unstratified and stratified by ASRM disease stage(*24*) (I-II and III-IV) and by menstrual cycle phase (proliferative, early secretory, and mid secretory) and found 299 drugs that reversed the endometriosis disease signature(*23*). Our top candidate, fenoprofen, an uncommonly prescribed non-steroidal anti-inflammatory drug (NSAID) in the U.S., successfully alleviated vaginal hyperalgesia in a rat model of endometriosis(*23*).

In the current study, we selected two therapeutic candidates from our earlier drug repositioning work for further investigation -- the cholesterol-lowering drug simvastatin and the antimalarial agent primaquine, based on their strong reversal of the endometrial signature in cases versus controls and good safety profiles(*25, 26*). Statins and anti-malarial agents have previously been considered potential therapeutics for endometriosis(*27–40*). Herein, we provide further evidence for simvastatin and primaquine as potential endometriosis therapeutics. Specifically, we conducted behavioral studies using an established rat model of endometriosis to assess the effect of treatment with simvastatin and primaquine on pain endpoints. Furthermore, we performed RNA sequencing on the uteri and lesions from these rats and analyzed differentially expressed genes (DEGs) for treated vs untreated (control) tissues. We also conducted a retrospective multi-center cohort study using a database of clinical records representing approximately 9.8 million individuals and assessed the risk of endometriosis among women prescribed simvastatin compared to those who were not prescribed this drug.

## RESULTS

### Overview

An overview of our study is provided in Fig. 1A. We previously leveraged a transcriptomics-based computational drug repositioning pipeline(*17*) using gene expression signatures of endometriosis queried against the CMap database(*41*), and identified fenoprofen as the top drug among 299 unique therapeutic candidates(*23*) (Table S1). Fenoprofen alleviated vaginal hyperalgesia comparably to our positive control, ibuprofen, in our rat endometriosis model(*23*). Among the compounds of interest with the greatest endometriosis disease-signature reversal, we selected two therapeutic candidates for further evaluation in this current study. In our endometriosis animal study model, we assessed pain-associated behavior in animals who were treated with each compound and compared them to untreated animals. We then performed RNA sequencing of the uterui and endometriosis lesions to assess the gene expression from these tissues in the treated and untreated animals.

**Fig. 1.**
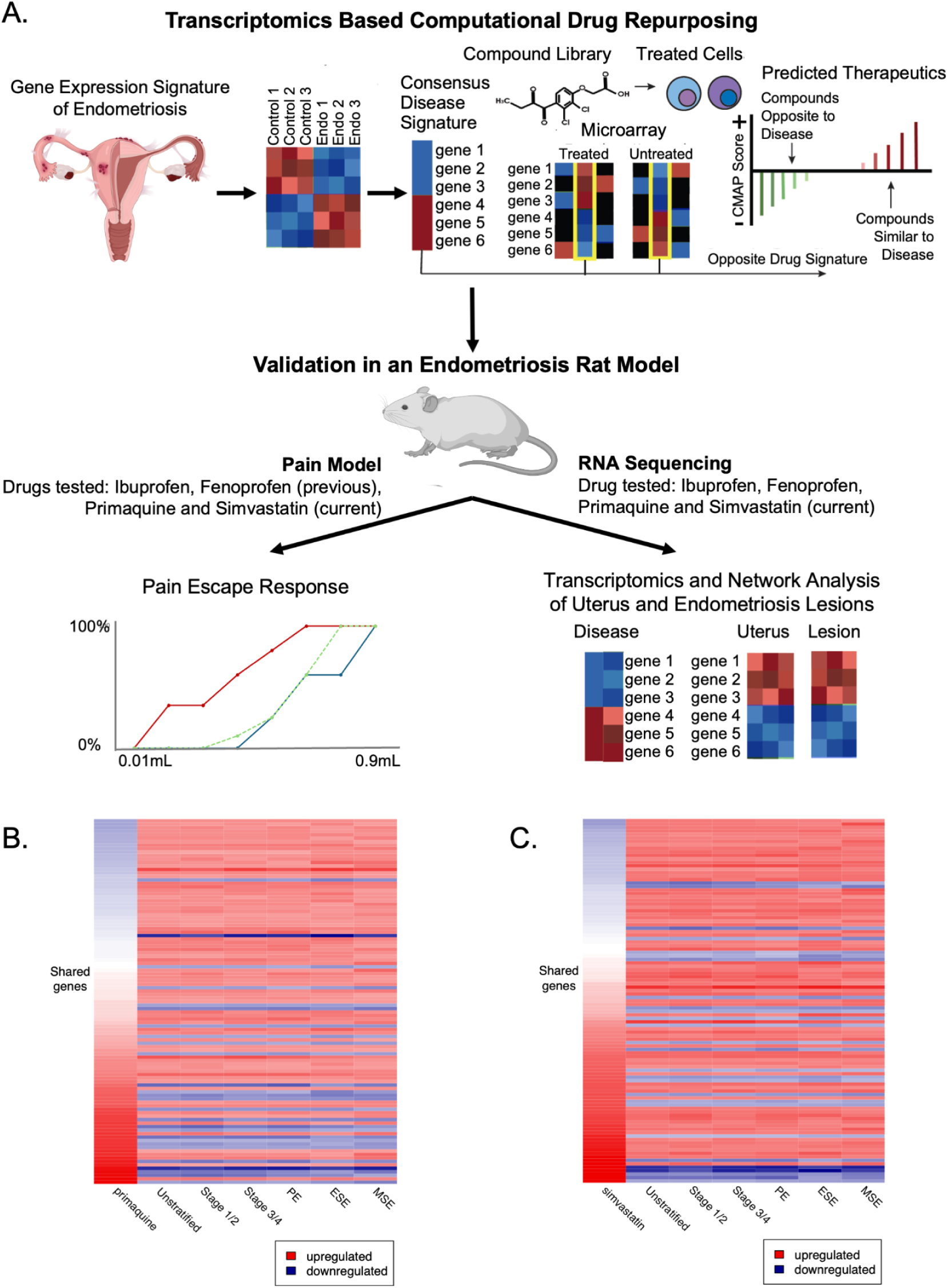
Study overview and disease signature reversal by primaquine and simvastatin. **(A)** Overview of the study that identified therapeutic candidates through a transcriptomics-based computational drug repositioning pipeline and then validated drug candidates in a rat model of endometriosis. Bulk RNA-sequencing was conducted of the uterus and endometriosis lesions from the rats, and differentially expressed genes and enriched pathways were analyzed. **(B,C)** Heatmap showing **(B)** primaquine and **(C)** simvastatin drug signature (far-left column of heatmaps) vs. unstratified and stratified (by stage and menstrual cycle phase) endometriosis disease signatures (remaining columns of heatmaps).

### Computational Drug Repositioning Pipeline Identifies Primaquine and Simvastatin as Therapeutic Candidates for Endometriosis

Among 299 drugs identified from the unstratified and ASRM disease stage- and cycle phase-stratified signatures, the cholesterol-lowering drug simvastatin and antimalarial agent primaquine were endometriosis therapeutic candidates selected for further study based on criteria including their strong reversal scores, safety profiles, and availability. When visualizing the gene expression of the six input endometriosis signatures (i.e., unstratified, and stratified by stage (I-II or III-IV) and phase (proliferative, early secretory, mid-secretory)) in comparison to the signature of simvastatin from CMap (Fig. 1B), the overall reversal pattern can be observed. A similar overall pattern of reversal can be seen when visualizing the gene expression of the six input endometriosis signatures and primaquine (Fig. 1C).

### Primaquine and Simvastatin Attenuate Escape Responses in a Rat Model of Endometriosis

The effect of our candidate drugs on endometriosis-associated vaginal hyperalgesia, a surrogate marker for endometriosis-related pain, was assessed in a rat model of endometriosis.

#### Vehicle

Among rats that received endometriosis (“endo”) surgery and vehicle treatment (n = 6), escape responses were significantly increased during the post-endo period compared to the baseline period, when volumes of 0.15, 0.30, 0.40, 0.55, 0.70, and 0.80 mL of water were delivered intra-vaginally (Mann–Whitney U test, Bonferroni-corrected p-value threshold of 0.05. Fig. 2B and Fig. S1). During the post-treatment period for the vehicle treatment group, similar to the post-endo period, escape responses were significantly increased relative to the baseline period when volumes of 0.15, 0.30, 0.40, 0.55, 0.70, and 0.80 mL of water were delivered (Mann–Whitney U test, Bonferroni-corrected p-value threshold of 0.05. Fig. 2B and Fig. S1). In the vehicle group, no statistically significant differences were found between the post-endo period and the post-treatment period escape responses for any volume of water delivered (Mann Whitney U test, Bonferroni-corrected p-value threshold of 0.05. Fig. 2B and Fig. S1).

**Fig. 2.**
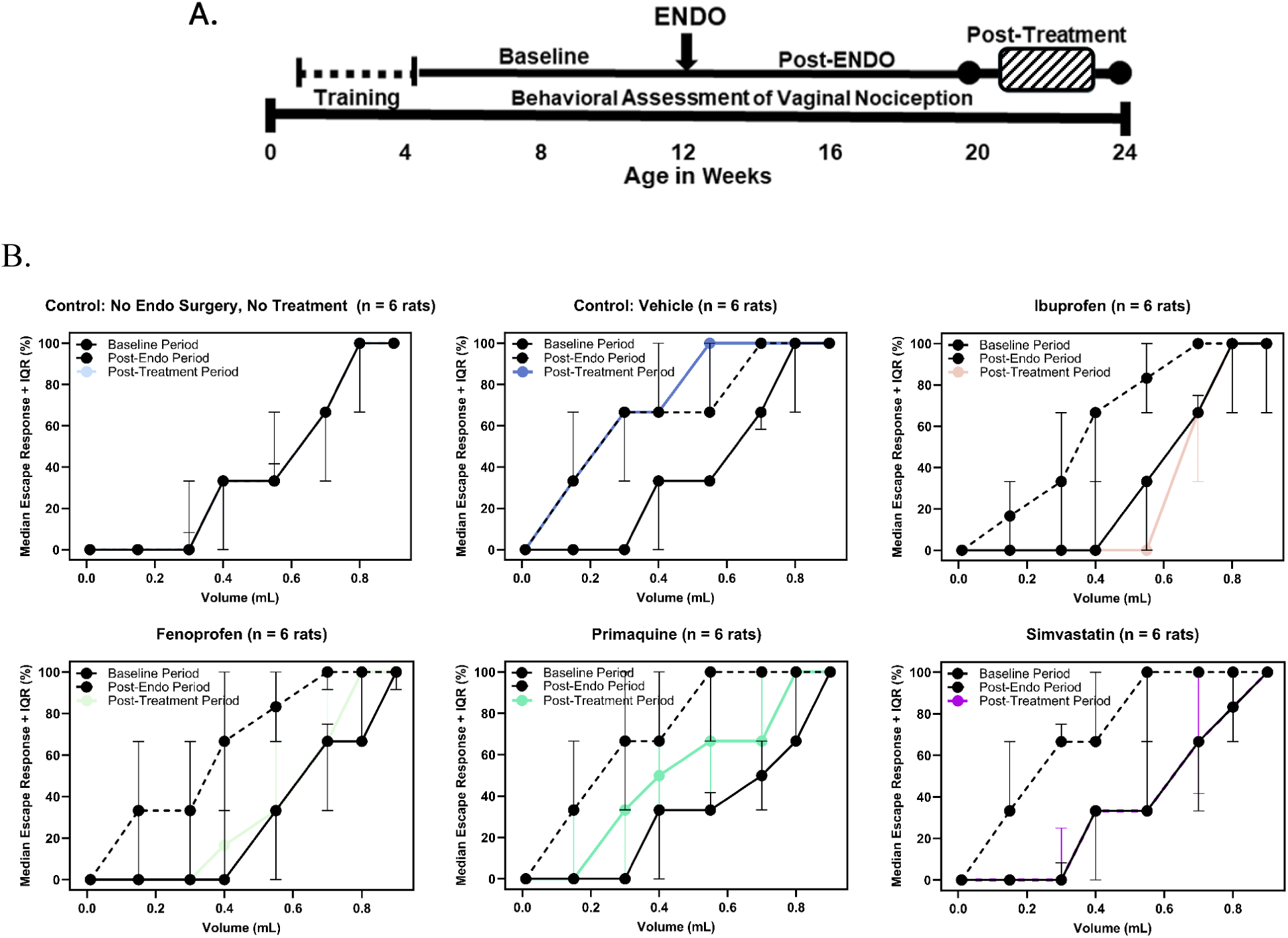
Endometriosis pain-associated escape behaviors in a rat model. **(A)** Female Sprague Dawley rats (n=18) were trained over 4 weeks to perform an escape response to terminate a noxious vaginal stimulus (water-filled balloon). Vaginal nociception was assessed as % escape response to varying balloon volumes in 1-hour sessions, 3 times/week, over 24 weeks. Each session included 8 volumes (0.01–0.90 mL), each tested 3 times in randomized, blinded order. Responses to none of the trials were counted as 0% escape response, 1 out of 3 trials as 33% escape response, 2 out of 3 trials as 67% escape response, and all 3 trials of a volume were counted as 100% escape response. Nociception was measured at Baseline (for 8 weeks), after endometriosis induction (Post-Endo, for 8 weeks), and during a 4-week treatment period. **(B)** Animal model median escape response (%) with interquartile range (IQR; error bars)(y-axis) for each delivered volume (0.01, 0.15,0.30, 0.40, 0.55, 0.70, 0.80, and 0.90 mL)(x-axis) during the baseline, post-endo surgery, and post-treatment periods for the 6 animal study groups; Control No endo Surgery; Vehicle; Ibuprofen; Fenoprofen; Primaquine; and Simvastatin. (Note: Control No endo Surgery, Ibuprofen, and Fenoprofen escape responses results are from previous work(*23*).)

#### Primaquine

Among rats that received endo surgery and primaquine treatment (40 mg/kg/day, p.o.) (n = 6), escape responses were significantly increased during the post-endo period compared to the baseline period, when volumes of 0.15, 0.30, 0.40, 0.55, 0.70, and 0.80 mL of water were delivered (Mann Whitney U test, Bonferroni-corrected p-value threshold of 0.05. Fig. 2C and Fig. S1). During the post-treatment period, escape responses were significantly decreased compared to the post-endo period, when volumes of 0.15, 0.30, 0.40, 0.55, and 0.70 mL of water were delivered (Mann Whitney U test, Bonferroni-corrected p-value threshold of 0.05. Fig. 2C and Fig. S1). Relative to the baseline period, post-treatment escape responses were significantly increased, when volumes of 0.30, 0.40, 0.55, and 0.70 mL of water were delivered (Mann Whitney U test, Bonferroni-corrected p-value threshold of 0.05. Fig. 2C and Fig. S1).

#### Simvastatin

Among rats that received endo surgery and simvastatin treatment (40 mg/kg/day, p.o.) (n = 6), escape responses were significantly increased during the post-endo period compared to the baseline period, when volumes of 0.15, 0.30, 0.40, 0.55, 0.70, and 0.80 mL of water were delivered (Mann Whitney U test, Bonferroni-corrected p-value threshold of 0.05. Fig. 2D and Fig. S1). During the post-treatment period, escape responses were significantly decreased compared to the post-endo period when volumes of 0.15, 0.30, 0.40, 0.55, 0.70, and 0.80 mL of water were delivered (Mann Whitney U test, Bonferroni-corrected p-value threshold of 0.05. Fig. 2D and Fig. S1). In the simvastatin group, no statistically significant differences were found between the baseline period and the post-treatment period escape responses for any volume of water delivered (Mann Whitney U test, Bonferroni-corrected p-value threshold of 0.05. Fig. 2D and Fig. S1).

#### Comparison Across Groups

Overall, the pattern of escape responses for primaquine (“PRIMA”) and simvastatin (“SIMVA”) are similar to those previously shared for fenoprofen-treated (“FEN”) and ibuprofen-treated (“IBU”, positive control) rats(*23*) (Fig. 2B and Fig. S1).

During the baseline period, there was a significant difference in escape responses between Control No endo Surgery (“CNS”) and Vehicle (“VEH”), and CNS and PRIMA at 0.55mL, and between PRIMA and IBU at 0.7mL, otherwise no other significant differences were identified between the 6 animal study groups (Fig. 3).

**Fig. 3.**
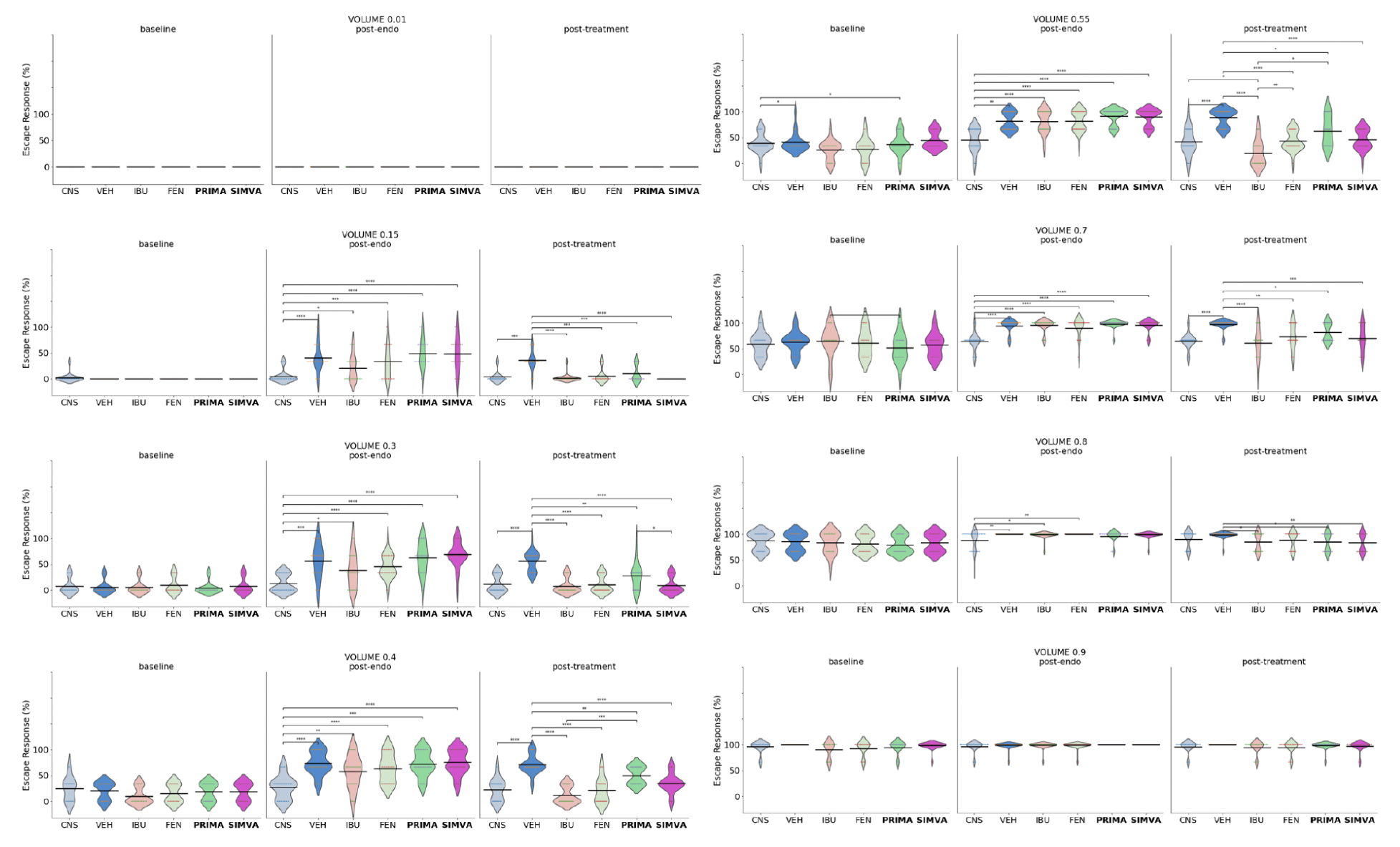
Strip and Violin plots with Medians and Significance Bars of Escape Responses to vaginal balloon distention at volumes of 0.01, 0.15, 0.30, 0.40, 0.55, 0.70, 0.80, and 0.90 mL for at baseline (Baseline), post-endometriosis (Post-Endo), and post-treatment (Post-Treatment) periods in a rat endometriosis model comparing 6 animal study groups — Control No Surgery (“CNS”), Vehicle (“VEH”), Ibuprofen (“IBU”), Fenoprofen (“FEN”), Primaquine (“PRIMA”), Simvastatin (“SIMVA”). Mann-Whitney test; Bonferonni adjusted p-values from Mann-Whitney U comparisons of groups. (Note: CNS, FEN, and IBU escape responses results are from previous work(*23*).) p-value annotation legend: ns: 5.00e-02 < p <= 1.00e+00 *: 1.00e-02 < p <= 5.00e-02 **: 1.00e-03 < p <= 1.00e-02 ***: 1.00e-04 < p <= 1.00e-03 ****: p <= 1.00e-04

During the post-endo period, there were significant differences between the negative control CNS and the 5 other groups at volumes 0.15, 0.3, 0.4, 0.55, and 0.7 mL; and between CNS and 3 other groups — VEH, IBU, and FEN — at 0.8mL No significant differences were identified between any of the 6 groups at 0.01 and 0.9mL (Fig. 3).

During the post-treatment period, there were significant differences in escape responses between the negative control VEH and the other 4 drug-treatment groups — IBU, FEN, PRIMA, and SIMVA — at volumes 0.15, 0.3, 0.4, 0.55, 0.7 and 0.8 mL; however, no significant differences between the groups were identified at 0.01 or 0.9 mL. There was a significant difference between CNS and IBU at 0.55 mL; otherwise, there were no significant differences in escape responses between CNS and the 4 drug-treatment groups — IBU, FEN, PRIMA, and SIMVA. Between the drug-treatment groups, there were significant differences in escape responses between PRIMA and SIMVA at 0.3 mL, PRIMA and positive control IBU at volumes 0.4 and 0.55 mL, and FEN and positive control IBU at volume 0.55 mL; otherwise no other significant differences were identified between positive control IBU, FEN, PRIMA, and SIMVA (Fig. 3).

### RNA-Seq Analysis of Treated Animals Confirms Reversal of Disease Signatures

To assess the impact of our candidate drugs on gene expression, bulk RNA sequencing was performed on the uterus and endometriosis-like “lesions” from rats (n = 6 per group per sample source).

#### Principal component analysis

Principal component analysis (PCA) revealed that samples from different treatment groups did not show distinct clustering (Fig. 4A). Instead, the sample source (i.e., uterus or lesion) was the primary factor contributing to the differences between samples (Fig. 4B). PCA was subsequently conducted within each sample source which still did not show distinct clustering based on treatment groups (Fig. S2).

**Fig. 4.**
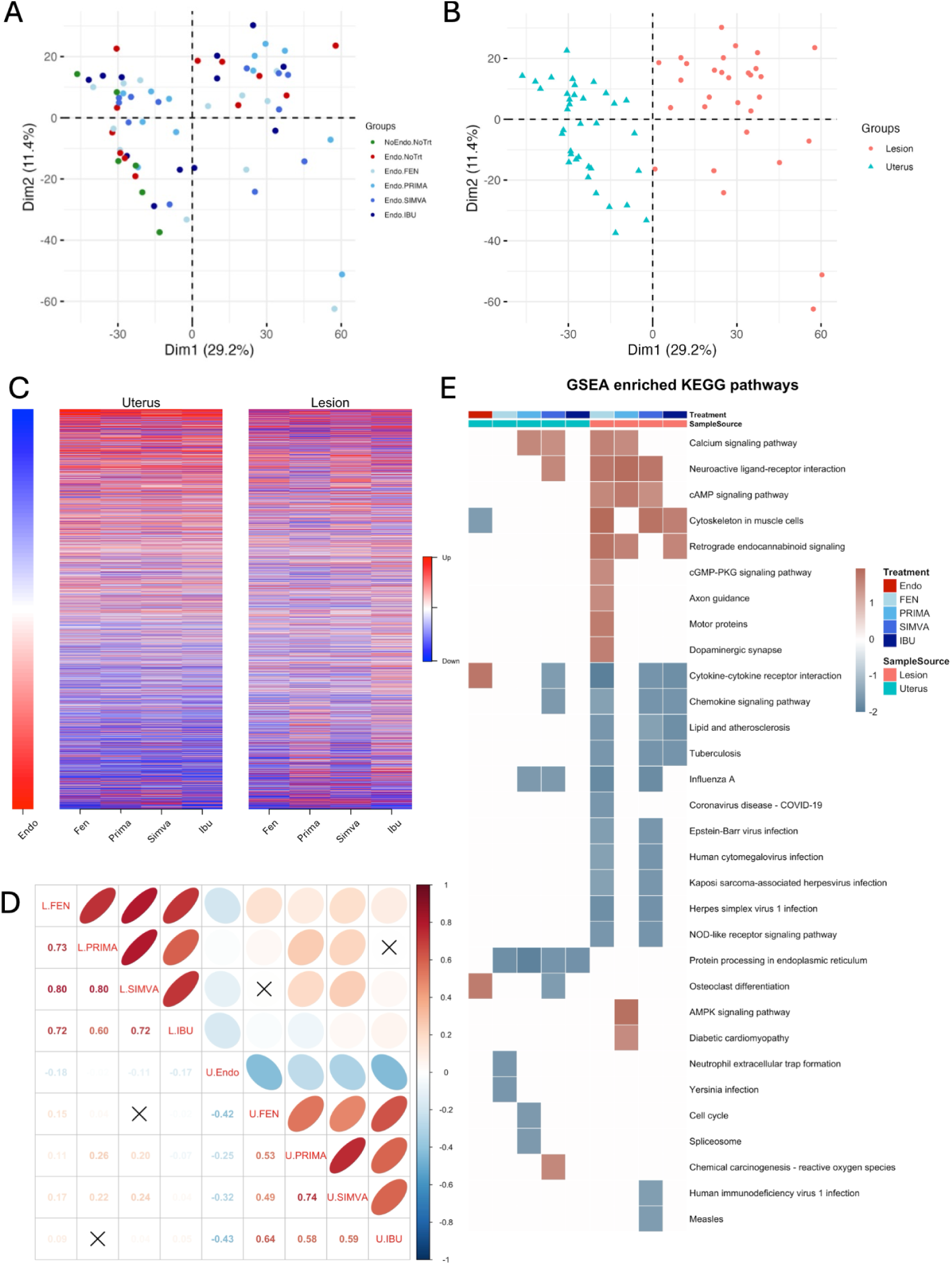
RNA-Seq analysis of treated animals. Principal component analysis (PCA) of all animal study samples, with each dot representing a sample, colored **(A)** by treatment group or **(B)** by sample source. **(C)** Heatmap of disease-associated (i.e., DEGs of endometriosis vs control) and treatment-associated (i.e., DEGs of treated vs control) gene signatures (Left: vehicle-treated uterus samples. Middle: drug-treated uterus samples. Right: lesion samples). Red indicates up-regulated genes and blue indicates downregulated genes. **(D)** Pearson correlation analysis between disease-induced (“U.Endo”) and drug-induced gene expression changes. Red indicates positive correlations and blue indicates negative correlations. **(E)** Gene Set Enrichment Analysis (GSEA) of KEGG pathways in disease and drug-treated groups. Red indicates up-regulated pathways and blue indicates downregulated pathways.

#### Differential gene expression

Differential gene expression analysis of the lesion and uterus samples between endometriosis rats with and without treatment revealed varying numbers of differentially expressed genes (DEGs) (Fig. S3 and Fig. S4, Tables S2-S10). Comparison of uterine samples from control rats and vehicle-treated rats that underwent endometriosis surgery revealed gene expression patterns associated with the disease (Fig. 4C, left). Further comparisons of the disease-associated gene expression patterns with those from drug-treated rats demonstrated that the four drugs --primaquine and simvastatin, as well as fenoprofen and positive control ibuprofen -- effectively reversed the gene expression profile in the uterus (Fig. 4C, middle), with significant negative correlations (Fig. 4D, bottom right). In the treated lesions, reversal of the gene expression profile associated with the disease was observed, particularly with fenoprofen, simvastatin, and ibuprofen (Fig. 4C, right), although compared with the treated uterine samples (Fig. 4C, middle), this reversal in the treated lesions appears less pronounced (Fig. 4C, right), and with only weak negative correlations (Fig. 4D, top left).

#### Gene set enrichment analysis

From gene set enrichment analysis, pathways enriched in the uterus before and after drug treatment were identified. *Cytokine-cytokine receptor interaction* and *osteoclast differentiation* were upregulated KEGG pathways in the uteri of endometriosis rats without drug treatment (i.e., vehicle), while *cytoskeleton in muscle cells* was a downregulated KEGG pathway. Simvastatin was able to reverse the *cytokine-cytokine receptor interaction* and *osteoclast differentiation* pathways in the uterus. Additionally, fenoprofen, simvastatin, and ibuprofen were able to reverse the *cytokine-cytokine receptor interaction* and *cytoskeleton in muscle cell pathways* in the lesion, although these effects were not strongly reflected at the overall gene expression level. (Fig. 4E).

#### Network analysis

To explore the similarities between the rat model and humans, the rat endometriosis gene signatures were compared to the human signatures reported in previous work(*23, 42*), resulting in the identification of 55 overlapping DEGs (Fig. 5A). A hypergeometric test on rat DEGs with human homologs confirmed their overrepresentation in human signatures. The expression profiles of these overlapping genes in the uterus and the lesion samples with and without drug treatment are visualized in Fig. 5B. Despite the foreseeable discrepancies in the expression profiles in humans and rats, several genes, such as Fos Proto-Oncogene (human *FOS*, rat *Fos)*, FosB Proto-Oncogene (human *FOSB*, rat Fosb), Interleukin 17C (Human *IL17C*, rat *Il17c)*, Nuclear Receptor Subfamily 4 Group A Member 1 (human *NR4A1*, rat *Nr4a1)*, Dual Specificity Phosphatase 1 (human *DUSP1*, rat *Dusp1)*, lymphocyte antigen 75 (human *LY75* or *DEC-205*, rat *Ly75*), and FRAS1 Related Extracellular Matrix 2 (human *FREM2*, rat *Frem2)* showed consistency across species. *FOS* and *FOSB*, members of the Fos gene family, encode Fos proteins which have previously been associated with endometriosis(*43, 44*). *NR4A1*(*45, 46*) and *LY75*(*47*) have also been linked with endometriosis, whereas *IL17C*, *DUSP1*, and *FREM2* are more novel for this condition. Notably, the four drugs reversed or partially reversed the expression of these genes in the uterus of endometriosis rats (Fig. 5B). To identify pathways associated with these 55 genes, we conducted a network analysis, revealing *cytokine-cytokine receptor interaction*, *IL-17 signaling*, and *MAPK signaling* as the primary pathways involved (Fig. 5C).

**Fig. 5.**
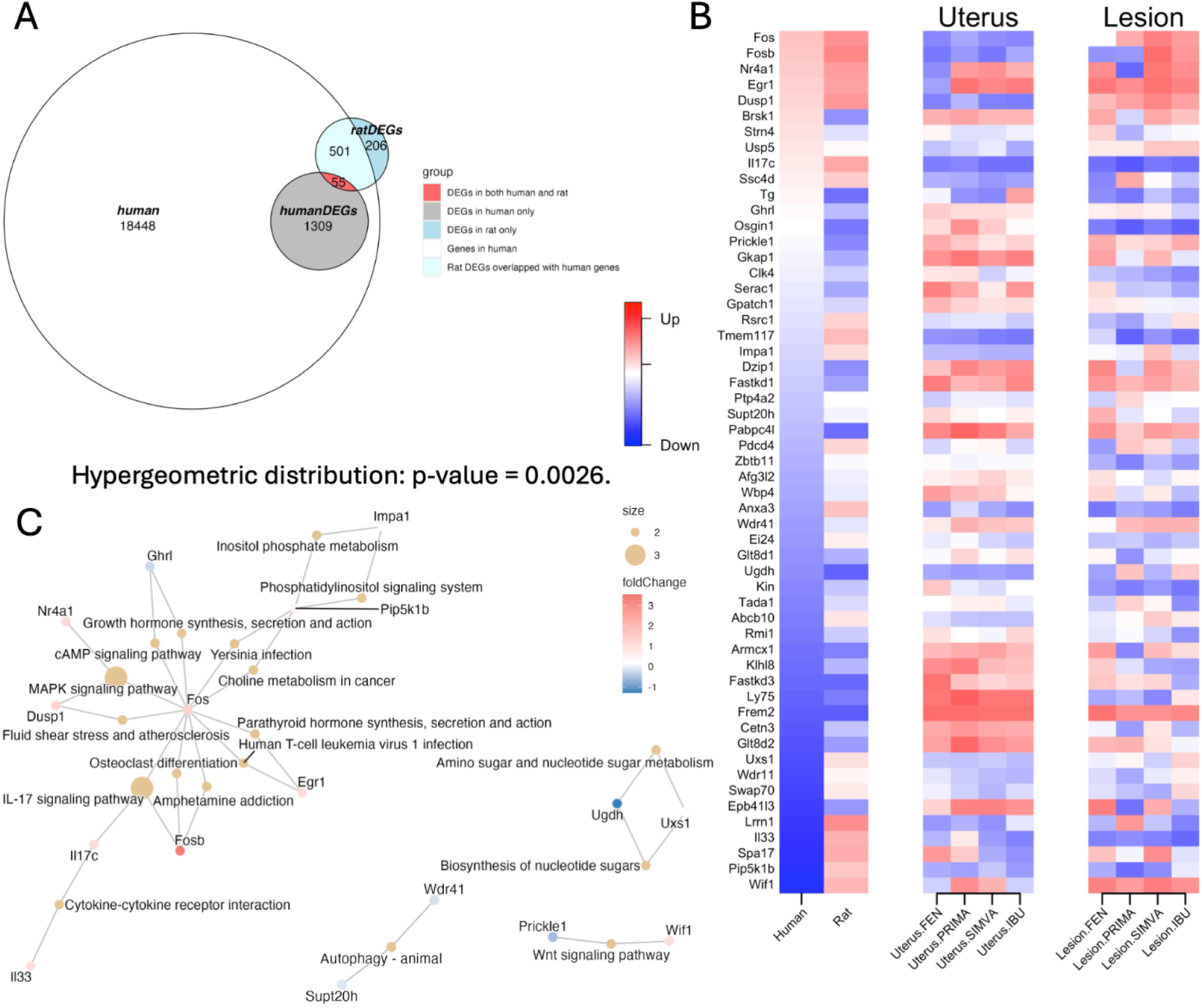
Correlation with human data. **(A)** Venn Diagram, **(B)** Heatmap, and **(C)** Network analysis to identify pathways associated with the 55 differentially expressed genes (DEGs) that overlap between our human and rat transcriptomics analysis, and their networks.

### Lower Endometriosis Risk with Simvastatin in Electronic Medical Records Analysis

Among approximately 9.8 million patients across six University of California health centers (January 1, 2012 - June 30, 2025) who were represented in the deidentified limited electronic medical records (EMR) dataset, 732 individuals had a prescription for primaquine and 265,754 individuals had a simvastatin prescription. After applying inclusion and exclusion criteria, the compound of interest (COI)-prescribed cohorts comprised 68 patients for primaquine (too few for statistically meaningful analysis) and 2,336 patients for simvastatin (Fig. S5).

Cohort characteristics and standardized mean differences (SMDs) of covariates between simvastatin-prescribed and control groups showed adequate balance after propensity score matching by demographics (age, race, ethnicity), UC healthcare site, number of visits, and medication indication (high cholesterol / hyperlipidemia), with the absolute value of SMD of less than 0.1 for all covariates after matching (Table 1, Table 2, Fig. S6).

**Table 1.** Cohort demographic characteristics before propensity score matching with absolute SMDs for patients prescribed simvastatin compared with control patients not treated with simvastatin.

| Characteristic | None | Simvastatin | SMD |
| --- | --- | --- | --- |
| **n** | 2,853,040 | 2,336 | - |
| **Year of Birth (mean (SD))** | 1974.95 (20.80) | 1981.68 (6.52) | 0.436 |
| **Age (median [IQR])** | 48.00 [33.00, 66.00] | 44.00 [40.00, 48.00] | 0.43 |
| **Race (%)** |  |  | 0.225 |
| - American Indian or Alaska Native (%) | 15,203 (0.5) | 21 (0.9) | - |
| - Asian (%) | 375,083 (13.1) | 263 (11.3) | - |
| - Black or African American (%) | 177,701 (6.2) | 204 (8.7) | - |
| - Native Hawaiian or Other Pacific Islander (%) | 18,412 (0.6) | 28 (1.2) | - |
| - Other Race (%) | 546,376 (19.2) | 608 (26.0) | - |
| - White (%) | 1,720,265 (60.3) | 1,212 (51.9) | - |
| **Ethnicity Hispanic or Latino (%)** | 603163 (21.1) | 782 (33.5) | 0.28 |
| **Location (%)** |  |  | 0.331 |
| - UC1 (%) | 451,026 (15.8) | 306 (13.1) | - |
| - UC2 (%) | 534,668 (18.7) | 631 (27.0) | - |
| - UC3 (%) | 361,474 (12.7) | 470 (20.1) | - |
| - UC4 (%) | 674,491 (23.6) | 413 (17.7) | - |
| - UC5 (%) | 817,636 (28.7) | 502 (21.5) | - |
| - UC6 (%) | 13,745 (0.5) | 14 (0.6) | - |
| **Number of visits (mean (SD))** | 31.82 (71.36) | 85.39 (140.29) | 0.481 |
| **High cholesterol / hyperlipidemia (%)** | 388,412 (13.6) | 1,189 (50.9) | 0.87 |
| **Endometriosis (%)** | 30,944 (1.1) | 67 (2.9) | 0.128 |

**Table 2.** Cohort demographic characteristics after propensity score matching with absolute SMDs for patients prescribed simvastatin compared with control patients not treated with simvastatin.

| Characteristic | None | Simvastatin | SMD |
| --- | --- | --- | --- |
| **n** | 9344 | 2,336 | - |
| **Year of Birth (mean (SD))** | 1981.67 (6.60) | 1981.68 (6.52) | 0.001 |
| **Age (median [IQR])** | 44.00 [40.00, 48.00] | 44.00 [40.00, 48.00] | <0.001 |
| **Race (%)** |  |  | 0.041 |
| - American Indian or Alaska Native (%) | 101 (1.1) | 21 (0.9) | - |
| - Asian (%) | 988 (10.6) | 263 (11.3) | - |
| - Black or African American (%) | 774 (8.3) | 204 (8.7) | - |
| - Native Hawaiian or Other Pacific Islander (%) | 132 (1.4) | 28 (1.2) | - |
| - Other Race (%) | 2519 (27.0) | 608 (26.0) | - |
| - White (%) | 4830 (51.7) | 1,212 (51.9) | - |
| **Ethnicity Hispanic or Latino (%)** | 3251 (34.8) | 782 (33.5) | 0.028 |
| **Location (%)** |  |  | 0.095 |
| - UC1 (%) | 1264 (13.5) | 306 (13.1) | - |
| - UC2 (%) | 2746 (29.4) | 631 (27.0) | - |
| - UC3 (%) | 1785 (19.1) | 470 (20.1) | - |
| - UC4 (%) | 1374 (14.7) | 413 (17.7) | - |
| - UC5 (%) | 2104 (22.5) | 502 (21.5) | - |
| - UC6 (%) | 71 (0.8) | 14 (0.6) | - |
| **Number of visits (mean (SD))** | 73.27 (127.14) | 85.39 (140.29) | 0.091 |
| **High cholesterol / hyperlipidemia (%)** | 4757 (50.9) | 1,189 (50.9) | <0.001 |
| **Endometriosis (%)** | 357 (3.8) | 67 (2.9) | 0.053 |

The rate of endometriosis among simvastatin-prescribed patients was 2.9% (67 of 2336) and among matched untreated control patients ranged from 3.8% (357 of 9344) to 4.1% (385 of 9344), with a 25% reduction in the relative risk (RR) of endometriosis (0.75 [95% CI, 0.58–0.97]; BH-corrected P-value = .03, with any reduction in the RR significant for 10 of 10 iterations)(Table 3).

**Table 3.**
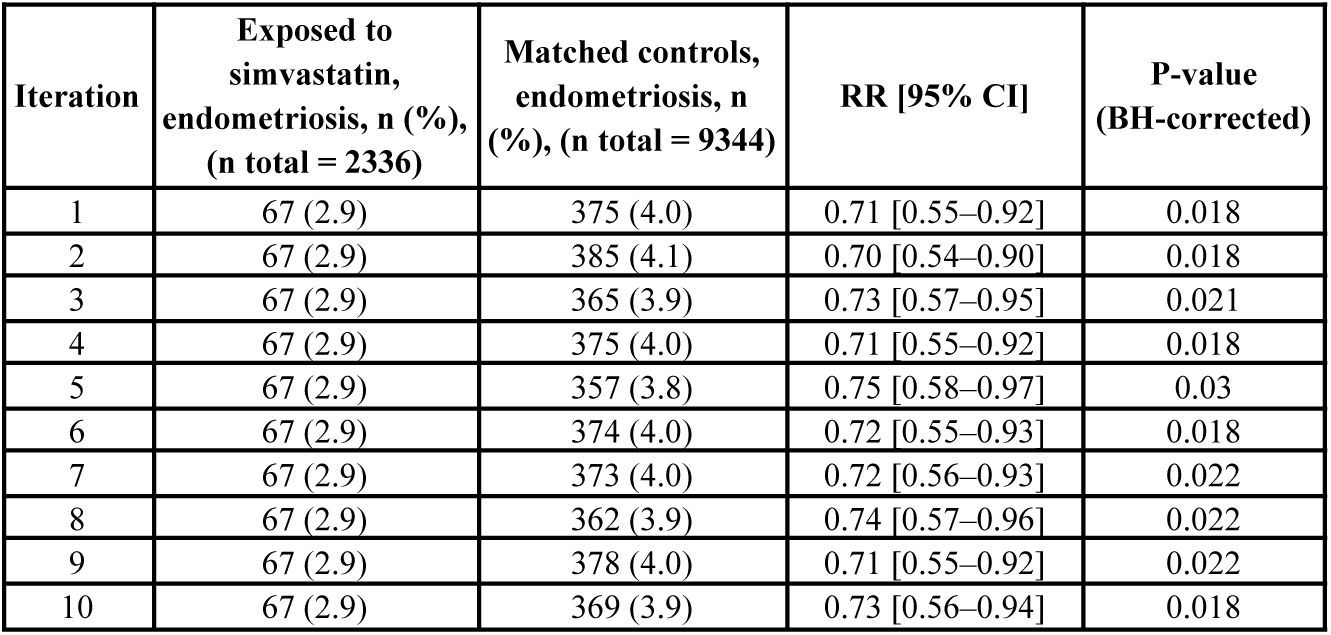
Relative risk of endometriosis in simvastatin-prescribed patients versus propensity score–matched unexposed controls (matched on age, race/ethnicity, UC health center, high cholesterol / hyperlipidemia, and number of visits).

| Iteration | Exposed to simvastatin, endometriosis, n (%), (n total = 2336) | Matched controls, endometriosis, n (%), (n total = 9344) | RR [95% CI] | P-value (BH-corrected) |
| --- | --- | --- | --- | --- |
| 1 | 67 (2.9) | 375 (4.0) | 0.71 [0.55–0.92] | 0.018 |
| 2 | 67 (2.9) | 385 (4.1) | 0.70 [0.54–0.90] | 0.018 |
| 3 | 67 (2.9) | 365 (3.9) | 0.73 [0.57–0.95] | 0.021 |
| 4 | 67 (2.9) | 375 (4.0) | 0.71 [0.55–0.92] | 0.018 |
| 5 | 67 (2.9) | 357 (3.8) | 0.75 [0.58–0.97] | 0.03 |
| 6 | 67 (2.9) | 374 (4.0) | 0.72 [0.55–0.93] | 0.018 |
| 7 | 67 (2.9) | 373 (4.0) | 0.72 [0.56–0.93] | 0.022 |
| 8 | 67 (2.9) | 362 (3.9) | 0.74 [0.57–0.96] | 0.022 |
| 9 | 67 (2.9) | 378 (4.0) | 0.71 [0.55–0.92] | 0.022 |
| 10 | 67 (2.9) | 369 (3.9) | 0.73 [0.56–0.94] | 0.018 |

## DISCUSSION

Effective therapeutics for endometriosis are in great need; however, developing new drugs for endometriosis is challenging due to the disease’s heterogeneity and complex symptoms, and the requirement of substantial resources and protracted timelines to bring new treatments to market(*13, 15*). Bioinformatics approaches can help identify existing drugs that can be repurposed to treat this condition(*16*). Our work leveraging publicly available endometrial disease and drug gene-expression signatures and a transcriptomics-based computational drug repositioning pipeline identified several FDA-approved drugs, including simvastatin and primaquine(*23*). While statins have been considered for some time as a potential treatment for endometriosis(*27–38, 43, 48, 49*), anti-malarial agents are more novel in this regard(*39, 40*). In vitro studies evaluating the effect of simvastatin on human endometrial stromal cell cultures have found that simvastatin inhibits cell growth in a concentration-dependent manner(*29–31*), and that lipid-soluble statins, including simvastatin, decrease stromal cell invasiveness(*32, 33*). A small prospective, randomized, double-blinded, controlled clinical trial (n = 60) comparing treatment with simvastatin versus a GnRH agonist in women who underwent laparoscopic surgery for pelvic endometriosis found significant reductions in dyspareunia, dysmenorrhea, and pelvic pain within both treatment groups over the 6-month postoperative period(*34*). No statistically significant differences were observed between the two treatment groups(*34*). A study of hydroxychloroquine found that treatment with this anti-malarial, in vitro, reduced endometrial and endometriotic cell survival and decreased the number of lesions in a mouse endometriosis model(*39*). Another study of the anti-malarial chloroquine and MK2206, an inhibitor of the serine/threonine protein kinase Akt (protein kinase B), found that combination therapy more effectively inhibited deep endometriotic stromal cell growth and reduced implant size in a mouse endometriosis model than either drug alone(*40*).

Our work leverages a murine endometriosis model, RNA sequencing, and real-world data to assess simvastatin and primaquine as potential therapeutics for endometriosis that could not only decrease the risk of disease but also alleviate endometriosis-related pain. While clinical records across the six healthcare centers included too few primaquine-prescribed patients for meaningful analysis, the dataset contained thousands prescribed simvastatin. Among 2,336 women who were prescribed simvastatin before age 40 and had no prior diagnosis of endometriosis, we observed a statistically significant 25% lower relative risk of endometriosis diagnosis compared with 9,334 propensity-score–matched controls who were never prescribed simvastatin. The findings from our analysis of real-world data are consistent with the possibility that simvastatin may reduce symptom severity or delay progression from subclinical disease to a clinically detectable stage.

We tested primaquine and simvastatin in a rat model of endometriosis, and found that treatment with each drug significantly alleviated vaginal hyperalgesia, a surrogate marker for endometriosis-related pain. Our findings with simvastatin were comparable to our prior findings with fenoprofen, the top drug candidate from our drug repositioning work, and with the positive control ibuprofen(*23*). There was a significant decrease in escape responses in the post-treatment period compared to the post-endo period (i.e., the period after the surgical induction of endometriosis). Furthermore, the degree to which simvastatin treatment attenuated the escape response was essentially the same as treatment with fenoprofen or ibuprofen, where there were no statistically significant differences between the baseline period and the post-treatment period(*23*). We found, however, that primaquine appeared less effective than ibuprofen, fenprofen, and simvastatin at attenuating the escape response. In endometriosis rats with no treatment, vaginal hyperalgesia was maintained, confirming that the reduction in hyperalgesia seen in the treatment groups was not a result of additional vaginal nociceptive testing following the establishment of endometriosis. Overall, the findings from our current animal study work support our drug candidates, especially simvastatin, as therapeutics that could be used for endometriosis-related pain.

Comparative analysis of gene expression in our animal model showed that treatment with primaquine, simvastatin, fenoprofen, and ibuprofen effectively reversed the disease-associated profile in the uteri, with significant negative correlations, which could reflect how the drugs are affecting aberrances within the uterus that are associated with endometriosis. Treatment with these drugs induced more modest yet directionally similar changes in the lesions, suggesting transcriptional responses to treatment that vary in degree by site. A plausible explanation for the limited transcriptomic reversal within lesions is that systemically administered drugs achieve lower and less sustained pharmacodynamic exposure in the ectopic implants than in eutopic uterus. In surgically induced rat endometriosis, implanted tissue fragments can develop hypoxia(*50*), inflammation(*51*), and fibrotic encapsulation(*52*), conditions that could substantially restrict penetration of medications including the compounds tested here. Therapeutic efficacy may depend on achieving sufficient exposure in lesions. Consequently, surgically induced lesions may require longer treatment, higher dosing, localized delivery, or agents that directly target stromal remodeling or fibrosis to produce stronger transcriptional effects.

Among 55 endometriosis disease-associated genes that were found in common between humans and the rat model, several including *Fos*, *Fosb*, *Nr4a1*, *and Ly75*, showed consistency in the direction of their expression across species, signaling the critical role of these genes in disease pathology, and a pattern of reversal of expression after treatment by one or more of our drugs. *FOS* and *FOSB* are members of the Fos gene family, which encode Fos proteins, components of the AP-1 transcription factor complex that regulates key cellular processes such as proliferation, migration, and transformation. *FOS* gene and Fos protein expression were found to be higher in endometriotic tissue and eutopic endometrium in women with endometriosis relative to the eutopic endometrium of patients without endometriosis(*44*). *FOS* has also been found to be upregulated during endometriosis establishment in a baboon endometriosis model(*43*). NR4A1 has been found to be overexpressed in endometriosis and NR4A1 antagonists inhibit the growth of endometriotic lesions(*45, 46*). Expression of *LY75*, involved in immune and inflammatory responses, has been found to be higher in patients with endometriosis compared to healthy control patients(*47*).

Gene set enrichment analysis identified pathways affected in our endometriosis animal model, including *cytokine-cytokine receptor interaction* and *osteoclast differentiation* which were upregulated in the uterus, and *cytoskeleton in muscle cells* which was downregulated, and that simvastatin reversed the *cytokine-cytokine receptor interaction* and *osteoclast differentiation* pathways in the uterus, and fenoprofen, simvastatin, and ibuprofen reversed the *cytokine-cytokine receptor interaction* and *cytoskeleton in muscle cell* pathways in the lesions. Furthermore, network analysis identified pathways associated with the endometriosis disease-associated genes common across the two species, revealing *cytokine-cytokine receptor interaction*, *IL-17 signaling*, and *MAPK signaling* as primary pathways involved. *Cytokine-cytokine receptor interaction* and *IL-17 signaling* play important roles in host immunity and inflammation, and have been considered among targets for immune-mediated inflammatory disorders such as rheumatoid arthritis and adenomyosis(*53–55*). *MAPK signaling* is crucial in regulating cellular processes such as proliferation, differentiation, development, transformation, and apoptosis(*56*). Interestingly, these pathways have also previously been identified as enriched in studies of ectopic endometrium surrounding ovarian cysts compared to eutopic endometrium of patients with endometriosis (*cytokine-cytokine receptor interaction*, *IL-17 signaling*, and *MAPK signaling*)(*57*) and human endometrial endothelial cells derived from eutopic endometrium of patients with and without endometriosis (*cytokine-cytokine receptor interaction*)(*58*). The enrichment of these pathways, particularly among conserved DEGs, suggests their critical role in endometriosis. Moreover, these pathways may indicate the mechanistic targets of candidate drugs. Endometriosis is characterized by the growth of endometrial tissue outside of the uterus and local and systemic inflammation; thus, targeting these pathways could provide therapeutic strategies to manage the disease and alleviate its symptoms.

### Limitations

Our study has several strengths, including robust and well-validated analgesic effects demonstrated in vivo, cross-species molecular analyses supporting reversal of key inflammatory pathways, and validation using real-world clinical data; however, it also has several limitations. Transcriptomics from endometrium, a tissue accessible from those with and without endometriosis, rather than endometriosis lesions was used to identify drug repositioning candidates. Matched control tissue for such lesions was not available in this study, and we are unable to generate robust differential expression signatures from rat lesion tissue alone to identify a disease signature. Eutopic and ectopic endometria of women with endometriosis have been found to exhibit shared molecular alterations that are absent in the eutopic endometria of women without the disease(*59*). Eutopic endometrium can offer important insights into disease mechanisms(*60*) and can serve as a practical foundation for therapeutic target discovery. Nevertheless, future work including signatures of the lesions themselves to query the drug data for therapeutic discovery would be valuable. Drugs not represented in the CMap dataset (e.g., ibuprofen and GnRH antagonists) would be missed by the drug repurposing pipeline. Similarly, drugs present in CMap but excluded during preprocessing due to inconsistent expression profiles also would not be identified(*20*). The drug repurposing pipeline focuses on identifying drugs that significantly reverse the overall endometriosis disease transcriptomic signature. This method does not necessarily consider whether the transcriptional effects of the drug are confined to the genes impacted by the disease. Therefore, a drug that induces extensive gene changes, including reversing the gene alterations caused by endometriosis, might be highlighted as a potential therapeutic option; however, such widespread gene changes may lead to unwanted side effects unrelated to the disease, which could limit the clinical usefulness of these drugs depending on the specificity and severity of the side effects and possible adverse pregnancy safety categories.

Moreover, reversal of disease signatures by our drug candidates was assessed only in the uteri, and effects on fertility were not directly evaluated; however, all rats continued to exhibit normal estrous cycles throughout the study, suggesting that gross reproductive function was not disrupted. While we and others have extensively utilized the CMap dataset for therapeutic discovery in various non-cancer conditions(*18, 19, 21–23*), it is important to note that the compounds in the CMap dataset were tested on cancer cell lines. The effects of these drugs on endometrial tissue and disease lesions would provide a more accurate assessment of their potential applications for treating endometriosis. Unfortunately, a large number of these compounds has not yet been tested on relevant endometrial tissues. However, as new datasets become available, we will incorporate them into our future drug discovery efforts for endometriosis. A further limitation is that our candidate therapeutics were evaluated in a rodent model. While this model reproduces several features of human endometriosis, rats neither menstruate nor develop the disease spontaneously. Menstruating nonhuman primates develop endometriosis naturally and therefore represent a more suitable translational model; however, their close genetic relatedness to humans entails distinct ethical and regulatory challenges. (*36, 43, 61*) Additionally, we evaluated a single dose level for both simvastatin and primaquine without formal PK or PD optimization or a dose response, which limits inferences about minimal effective doses and exposure response relationships. Also, our electronic medical records analysis was retrospective and therefore can establish association, not causation, between simvastatin exposure and reduced endometriosis risk. Medication histories and comorbidity data may be incomplete for some individuals, which could affect these findings. Prospective clinical trials are needed to determine therapeutic efficacy in endometriosis.

### Conclusions

Our work demonstrates that simvastatin and primaquine, therapeutic candidates identified by our transcriptomics-based drug repositioning computational pipeline, effectively alleviate pain-associated behavior in an animal model of endometriosis. Additionally, these drugs successfully reversed the disease signature at the gene expression level in this animal model. Simvastatin, in particular, shows promise as a potential therapeutic for endometriosis-related pain. Moreover, our analysis of real-world data indicated that women prescribed simvastatin experienced fewer endometriosis diagnoses than matched unexposed patients, consistent with the possibility that simvastatin may lessen symptom severity or delay progression from sub-clinical disease to a clinically detectable stage. The overlapping genes and pathways between our animal model and endometriosis patients offer insights into the biological processes associated with the disease and potential therapeutic targets. Also. our findings highlight the value of computational approaches in identifying therapeutic strategies for endometriosis.

## METHODS

### Computational Drug Repositioning

In previous work(*23*), our computational drug repositioning pipeline was applied to bulk transcriptomic data, which consisted of 105 samples from eutopic endometrial tissues of women with and without endometriosis. On the drug side, the Connectivity Map (CMap) dataset was used to obtain gene expression profiles from cell lines treated with existing small-molecule drugs. Through this approach, potential therapeutics were identified based on reversal of endometriosis gene expression signatures that were unstratified as well as stratified by ASRM disease stage (ASRM stages I/II and III/IV) and menstrual cycle phase (proliferative, early secretory, and mid-secretory phases) as described in Bunis et al.(*42*). As a proof of principle, the top therapeutic candidate fenoprofen, an NSAID infrequently prescribed for endometriosis, was validated in an animal model of endometriosis.

In this study, among the 299 unique drug candidates that were previously identified, those among the top-ranked (i.e., among the top 10%) were evaluated for further consideration, which included primaquine and simvastatin. Factors such as safety profiles, availability (i.e., on the World Health Organization list of essential medicines(*62*)), and ease of administration (e.g., does not require intravenous administration) were taken into consideration.

### Animal Studies

The primary aim of the animal study was to evaluate the effects of simvastatin and primaquine on pain-associated behavior in a rat model of endometriosis. At baseline, estrous cycles were monitored and nociceptive behavior was characterized in all animals. Endometriosis was then surgically induced, and rats received simvastatin, primaquine, or vehicle (negative control) for 4 weeks. Behavioral assessments of vaginal nociception were conducted after induction and again after treatment to determine treatment effects. Groups of six rats were assigned to each arm, and a within-subject design enabled comparisons across phases (baseline, post-induction, post-treatment). All training and testing occurred 3–8 hours after lights-on, three times per week on non-consecutive days.

#### Animal Subjects and Conditions

A total of 18 adult virgin female Sprague-Dawley rats were used in this study, each weighing between 175 and 225 grams at the onset of the study, and were sourced from Charles River Laboratory (Wilmington, MA; Raleigh, NC facility). Our study exclusively examined female rats because the disease modeled is only relevant in females. The animals were housed individually in standard rodent cages with access to water and chow ad libitum. They were kept under controlled conditions with a 12-hour light/dark cycle (lights on at 07:00), and the room temperature was maintained at ∼22°C. Estrous cycle stages were monitored and documented daily 2 hours after lights on via vaginal lavage.

#### Endometriosis Induction

Endometriosis was surgically induced in rats as described by Vernon and Wilson(*52*). Animals were anesthetized with a combination of ketamine (73 mg/kg) and xylazine (8.8 mg/kg), and a midline abdominal incision was made. A 1 cm segment of the left uterine horn with surrounding fat was excised and transferred to sterile saline. This tissue was cut into four 2x2 mm fragments and sutured onto the mesenteric arteries supplying the small intestine. After the surgical procedure, the incision was closed, and the rats were closely monitored during recovery. The operation had no complications and estrous cycles resumed within a few days.

#### Drug Administration

Rats were randomly assigned (n=6 per group) to receive simvastatin (40 mg/kg/day), primaquine (40 mg/kg/day), or vehicle by oral gavage once daily for 4 weeks after endometriosis induction. The 40 mg/kg/day dose for both agents was selected from prior rodent studies showing activity and tolerability in inflammatory and nociceptive models(*35, 63–67*). Vehicle animals received an equal volume of dosing vehicle without active drug. Behavioral testing and analysis were performed blinded to treatment assignment.

#### Assessment of Vaginal Nociception

Vaginal nociception was assessed using a behavioral test in which rats were exposed to vaginal distension via inflation of a latex balloon inserted into the vaginal canal. The primary endpoint was the percentage of successful nociceptive escape responses, with animals trained to break a light beam by extending their heads into a tube when they experienced discomfort from the distension. This procedure was repeated at eight different balloon volumes, three times each in random order, to characterize volume-dependent changes in nociceptive escape behavior.

#### Behavioral Testing Apparatus

The testing apparatus was a rectangular Plexiglas chamber equipped with a grid floor, which restricted the rat from turning around. A hollow tube containing light-emitting diodes and a photosensor extended from the front of the chamber. When the rat extended its head, it interrupted the light beam, resulting in the cessation of the stimulus and the provision of peanut butter as a reward by the experimenters. This action, referred to as an escape response, was used to measure behavioral responses to the vaginal distention.

An opening in the rear of the chamber allowed the balloon-tipped catheter to be connected to the computer-controlled and automated stimulus-delivery device. The latex balloon (10 mm long, 1.5 mm wide when uninflated) was lubricated with K-Y® jelly and inserted into the vaginal canal of the rat before each session. It was then inflated to different volumes through the computer-controlled pump to induce vaginal distention at 60-second intervals, with the pressure produced by each volume monitored via a small-volume Cobe pressure transducer and the escape response to each volume recorded.

#### Behavioral Training

Training began by acclimating the rats to the testing chamber for 10 minutes daily for 3–4 days, during which small amounts of peanut butter were provided on a wooden stick. The rats were then trained by using tail pinches with padded forceps, which prompted them to extend their head into the tube to break the light beam. Once the light beam was broken, the tail pinch was released to reinforce the “escape response,” and the rats were rewarded with peanut butter if they successfully interrupted the light beam to escape the tail pinch. This process was repeated for 10 tail pinches at 1-minute intervals over 3 training sessions each week on non-consecutive days. Training was completed (>80% escape behavior) in 4–8 sessions.

The rats were next trained to make identical escape responses to deflate vaginal distention stimuli. These sessions were run 3 times/week on non-consecutive days for a total of 3–5 sessions. Ten large distention volumes (0.80 ml – 1.0 ml, inflation rate 1 ml/s) were delivered for a maximum of 15 s at 1-min intervals. All rats showed some behavioral response to these stimuli, which allowed the experimenter to use deflation of the vaginal balloon to shape the rat’s escape responses. All rats learned the escape response within 2–4 sessions. Once trained, testing sessions began.

#### Experimental Procedure

Behavioral testing was conducted across three phases: baseline (prior to endometriosis induction), post-endo (after endometriosis induction but before treatment), and post-treatment (after drug/control administration). In each phase, rats underwent a series of 24 trials (8 different balloon inflation volumes delivered 3 times each). Each trial involved inflating the balloon at a constant rate of 1 mL/s and maintaining the inflation for up to 15 seconds unless an escape response occurred. The balloon was then deflated at 0.5 mL/s if an escape response occurred or if 15 seconds elapsed. The percentage of successful escape responses was recorded for analysis.

#### Statistical Analysis

For each rat group (VEH: with endo surgery, vehicle treatment (vehicle control) (n=6); (SIMVA: with endo surgery, treated with simvastatin (n=6); PRIMA: with endo surgery, treated with primaquine (n=6), a non-parametric test was performed to compare within groups, the escape responses of the 3 testing periods: baseline (24 data points/rat), post-endo surgery (64 data points/rat), and post-treatment (32 data points/rat). Mann Whitney U (MWU) tests were performed to compare the escape responses of (a) the baseline period and the post-endo period, (b) the post-endo period and the post-treatment period, and (c) the baseline period and the post-treatment period at each volume of water (0.0, 0.15, 0.30, 0.40, 0.55, 0.70, 0.80, and 0.90 mL) delivered to the balloon placed within the mid-vaginal canal of the rats. Moreover, Mann Whitney U (MWU) tests were performed to compare between groups, the escape responses of (a) VEH vs. SIMVA, (b) VEH vs. PRIMA, and (c) SIMVA vs. PRIMA for each of the three condition periods (the baseline period, the post-endo period, and post-treatment period) at each volume of water (0.01, 0.15, 0.30, 0.40, 0.55, 0.70, 0.80, and 0.90 mL) delivered to the balloon placed within the mid-vaginal canal of the rats. Escape responses reflect the percentage of times rats extended their head into the tube to interrupt the light beam to terminate the stimulus (balloon deflates). A significance threshold of 0.05 was applied to Bonferroni–corrected p-values. Data are presented as median and interquartile range (IQR).

### RNA Seq

#### Experimental Procedure

RNAs were extracted using miRNeasy mini Kit (Qiagen), and 1.5ug of total RNA was used for library construction. The libraries for RNA sequencing were prepared using the Illumina Stranded Total RNA Prep with Ribo-Zero^TM^ Plus kit. We sequenced libraries using Novaseq X-Plus (Illumina), paired-end, 100 reads up to 50M paired reads per sample.

#### Statistical Analysis

Adapter sequences in the raw read files were trimmed using cutadapt (v4.9) with an error rate of 0.1, minimum overlap of 3 bps, minimum 3’ base quality score of 25, and minimum length after trimming of 55 bps. We also set to remove poly-A tails and ploy-G with lengths over 20 bps. The trimmed reads were mapped to Rattus Norvegicus reference genome GRCr8 with exon and splice site information using Hisat2 (v2.2.0). The SAM files were converted to BAM files and sorted using samtools (v1.21). The gene counts were summarized by featureCounts (v2.0.6).

For the dimensionality reduction, gene counts were log_2_ transformed and the top 3000 most variable genes based on their standard deviation were used to perform principal component analysis (PCA).

The differential gene expression analysis comparing treatment groups was performed using the edgeR package (v4.4.1) in R 4.4.2. We first excluded genes that were uncharacterized from further analysis. Low-expressed genes were dropped using the filterByExpr() function based on the treatment factors. After filtering, we recalculated the library sizes. To account for sample-specific effects, we applied the trimmed mean of M-values (TMM) normalization. We then estimated the dispersions for all genes and fitted a negative binomial generalized log-linear model to the gene counts. This model included an interaction term between the sample source and treatment, with the RNA Integrity Number (RIN) included as a covariate. The differential expression of genes between treatment groups within each sample source was tested with genewise quasi F-tests for the coefficient contrasts of the model. P-values were subjected to the Benjamini-Hochberg (BH) method to control the false positive rate. A Benjamini-Hochberg (BH)-adjusted p-value threshold of 0.05 was used to determine significance. Gene set enrichment analysis and over-representation analysis were performed using the clusterProfiler package (version 4.14.4). Pearson correlation coefficients were calculated to assess gene expression correlations between treatment groups. The overlap between human disease signatures and rat differentially expressed genes was evaluated using the hypergeometric test.

### Electronic Medical Records

#### Cohort Definition

A HIPAA-compliant limited dataset of electronic medical records (January 1, 2012 - May 30, 2025) representing patients across six University of California health systems (UC Davis, UC Irvine, UCLA, UC Riverside, UC San Diego, UCSF) was accessed via the University of California Data Discovery Platform (UCDDP) and analyzed in August 2025. The number of individuals who were ever prescribed primaquine or simvastatin was assessed, and investigation of the compound of interest (COI) proceeded if there was sufficient power for downstream analysis. Our analysis compared endometriosis diagnoses among individuals with a prescription for the COI to a propensity-score–matched cohort without any such prescription. Conditions identified by using International Classification of Diseases, Ninth Revision, Clinical Modification (ICD-9-CM), and International Statistical Classification of Diseases, Tenth Revision, Clinical Modification (ICD-10-CM), diagnosis codes include endometriosis (617.X, N80.X) and high cholesterol/hyperlipidemia (272.0X, 277.1X, 277.2X, 277.3X, 277.4X, E78.0X, E78.1X, E78.2X, E78.3X, E78.4X, E78.5X).

Patients who were of female sex were included. Among patients with a prescription for the COI, they were included only if their first COI prescription occurred at age 40 or younger and at least five years had passed since that first COI prescription date. Individuals who were younger than 12 years old, missing information for sex, race, ethnicity, UC healthcare site, or number of visits, or had an endometriosis diagnosis prior to their first COI prescription were excluded.

#### Statistical Analysis

Propensity score matching was conducted with the MatchIt package (R, v4.7.2) using nearest-neighbor matching at a 1:4 treated-to-control ratio. Propensity scores were estimated via logistic regression of treatment on baseline demographics (age, race, ethnicity), UC healthcare site, number of visits, and medication indication. Covariate balance was assessed with balance plots of the standardized mean differences for covariates before and after propensity-score matching.

The matching procedure was repeated 10 times to address uncertainty from tied matches. Each iteration included all treated patients and a re-sampled, propensity-matched control subset. For each iteration, the relative risk of endometriosis diagnosis (with 95% confidence intervals) was estimated and results summarized across iterations to evaluate robustness. Multiple hypothesis testing was controlled using the Benjamini–Hochberg procedure with an adjusted significance threshold of 0.05.

### Study Approval

The animal study and procedures were approved by the Emory University Institutional Animal Care and Use Committee (IACUC) as protocol #2021000201. All laboratory animal experimentation adhered to the NIH Guide for the Care and Use of Laboratory Animals. The electronic medical records analysis were approved by the University of California, San Francisco Institutional Review Board (IRB) as protocol #22-37954.

### Data availability

The animal study RNA seq data are available through the Gene Expression Omnibus (GEO), accession ID GSE296883 (https://www.ncbi.nlm.nih.gov/geo/query/acc.cgi?acc=GSE296883). Data that support the findings of this study were obtained from the University of California Data Discovery Platform (UCDDP), a HIPAA limited data set which contains patient data from all 6 UC Health academic medical centers (Davis, Irvine, Los Angeles, Riverside, San Diego, and San Francisco). Due to privacy and confidentiality restrictions, the data are not publicly available. Minimal de-identified aggregate data in the form of tables are available from the corresponding author on reasonable request and subject to corresponding author’s institutional approval. UC researchers may obtain access after completing an initial analysis within their campus health system and securing approval for UC-wide extension. Additional instructions are available through the UC Health Center for Data-Driven Insights and Innovation (https://www.ucop.edu/uc-health/departments/center-for-data-driven-insights-and-innovations-cd i2.html).

## ACKNOWLEDGMENTS

**General:** The authors acknowledge the use of resources developed and supported by the UCSF Bakar Computational Health Sciences Institute Information Commons team, and thank members of this team for technical support. The authors also thank the Center for Data-driven Insights and Innovation at UC Health, and the members of their team for technical support.

## Funding

NIH NICHD P01 HD106414 (T.T.O., A.B., J.C.I., B.G., D.K.S., L.C.G., S.L.M., M.S.)

NIH NICHD P50 HD055764 (A.B., J.C.I., L.C.G., M.S.)

NIH NICHD R00 HD093858 (S.L.M)

March of Dimes Prematurity Research Center at UCSF (T.T.O., M.S.)

March of Dimes Prematurity Research Center at Stanford University (B.G., D.K.S.)

Stanford Maternal and Child Health Research Institute (B.G., D.K.S.)

Emory Integrated Genomics Core (EIGC) (RRID:SCR_023529), which is subsidized by the Emory University School of Medicine and is one of the Emory Integrated Core Facilities. Georgia Clinical & Translational Science Alliance of the National Institutes of Health under Award Number UL1TR002378.

The content is solely the responsibility of the authors and does not necessarily reflect the official views of the National Institutes of Health.

## Author Contributions

Conceptualization: TTO, XT, LCG, SLM, MS.

Methodology: TTO, XT, EA, AG, AL, SLM.

Investigation: TTO, XT, EA, AG, AL, SLM.

Visualization: TTO, XT, FA, SLM.

Funding acquisition: BG, DKS, LCG, SLM, MS

Supervision: BG, DKS, LCG, SLM, MS.

Writing – original draft: TTO, XT, EA, AG, FA, SLM.

Writing – review & editing: TTO, XT, EA, AG, FA, DJB, KE, JE, MD, BG, DKS, LCG, SLM, MS.

## Data and Materials

Data that support findings of this study were obtained from the University of California Data Discovery Platform (UCDDP), a HIPAA limited data set which contains patient data from all 6 UC Health academic medical centers (Davis, Irvine, Los Angeles, Riverside, San Diego, and San Francisco). Due to privacy and confidentiality restrictions, the data are not publicly available. Minimal de-identified aggregate data in the form of tables are available from the corresponding author on reasonable request and subject to corresponding author’s institutional approval.

## Competing Interests

A.B., D.B., T.T.O., L.C.G., and M.S. have a patent related to this work. L.C.G. is a paid consultant to Gensyta Pharma, Celmatix, and NextGen Jane. M.S. is an advisor to Wellcome Leap. The remaining authors declare no competing interests.

## LIST OF SUPPLEMENTARY FIGURES AND TABLES

**Fig. S1: Endometriosis animal model escape responses.**

**Fig. S2. Principal component analysis (PCA) for each sample source.**

**Fig. S3. Volcano plots of individual differentially expressed genes (DEGs) in lesion samples by treatment group.**

**Fig. S4. Volcano plots of individual differentially expressed genes (DEGs) in uterus samples by treatment group.**

**Fig. S5. Overview of the Analysis for Simvastatin in Electronic Medical Records**

**Fig. S6. Balance plot of the absolute standardized mean difference (SMD) of covariates before PS-matching and after PS-matching.**

**Table S2. Differentially expressed genes (DEGs) in lesion samples in FEN treatment group.**

**Table S3. Differentially expressed genes (DEGs) in lesion samples in IBU treatment group**

**Table S4. Differentially expressed genes (DEGs) in lesion samples in PRIMA treatment group.**

**Table S5. Differentially expressed genes (DEGs) in lesion samples in SIMVA treatment group.**

**Table S6. Differentially expressed genes (DEGs) in uterus samples in VEH treatment group.**

**Table S7. Differentially expressed genes (DEGs) in uterus samples in FEN treatment group.**

**Table S8. Differentially expressed genes (DEGs) in uterus samples in IBU treatment group.**

**Table S9. Differentially expressed genes (DEGs) in uterus samples in PRIMA treatment group.**

**Table S10. Differentially expressed genes (DEGs) in uterus samples in SIMVA treatment group.**

## SUPPLEMENTARY FIGURES

**Fig. S1:**
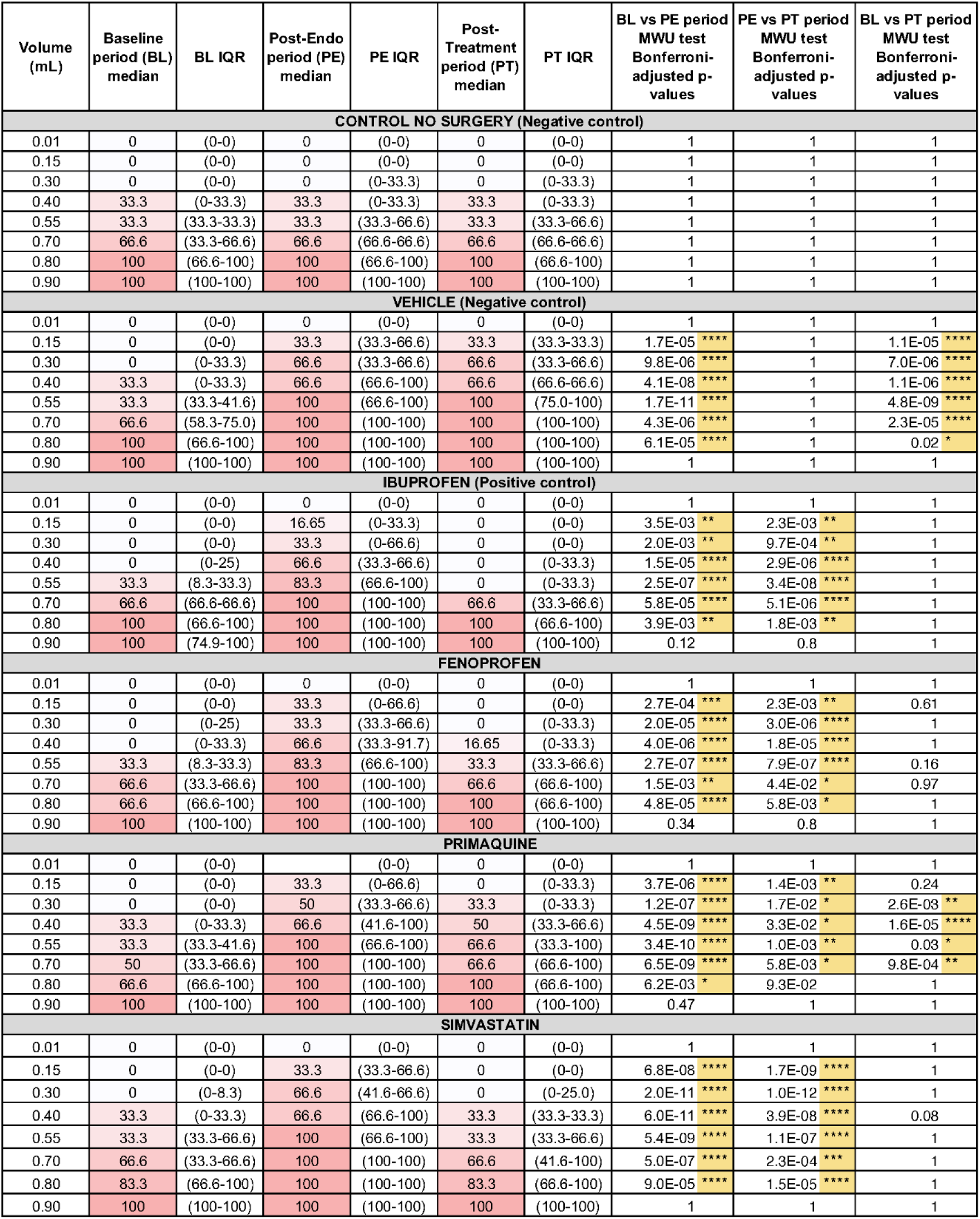
Endometriosis animal model escape responses. Median escape response with interquartile range (IQR) for each delivered volume (0.01, 0.15,0.30, 0.40, 0.55, 0.70, 0.80, and 0.90 mL) during the baseline, post-endo surgery, and post-treatment periods, with Bonferroni-corrected p-values from Mann-Whitney U test for baseline period vs post-endo surgery period, post-endo surgery vs post-treatment period, and baseline period vs post-treatment period (* denotes <0.05, ** denotes <0.005, *** denotes <0.0005, and **** denotes <0.0001). Endo rats (n=6/group) vaginal nociception assessed. Treated with vehicle (negative control), primaquine, simvastatin, fenoprofen, or ibuprofen (positive control). Note: Fenoprofen and ibuprofen data are from prior work(*23*)

**Fig. S2.**
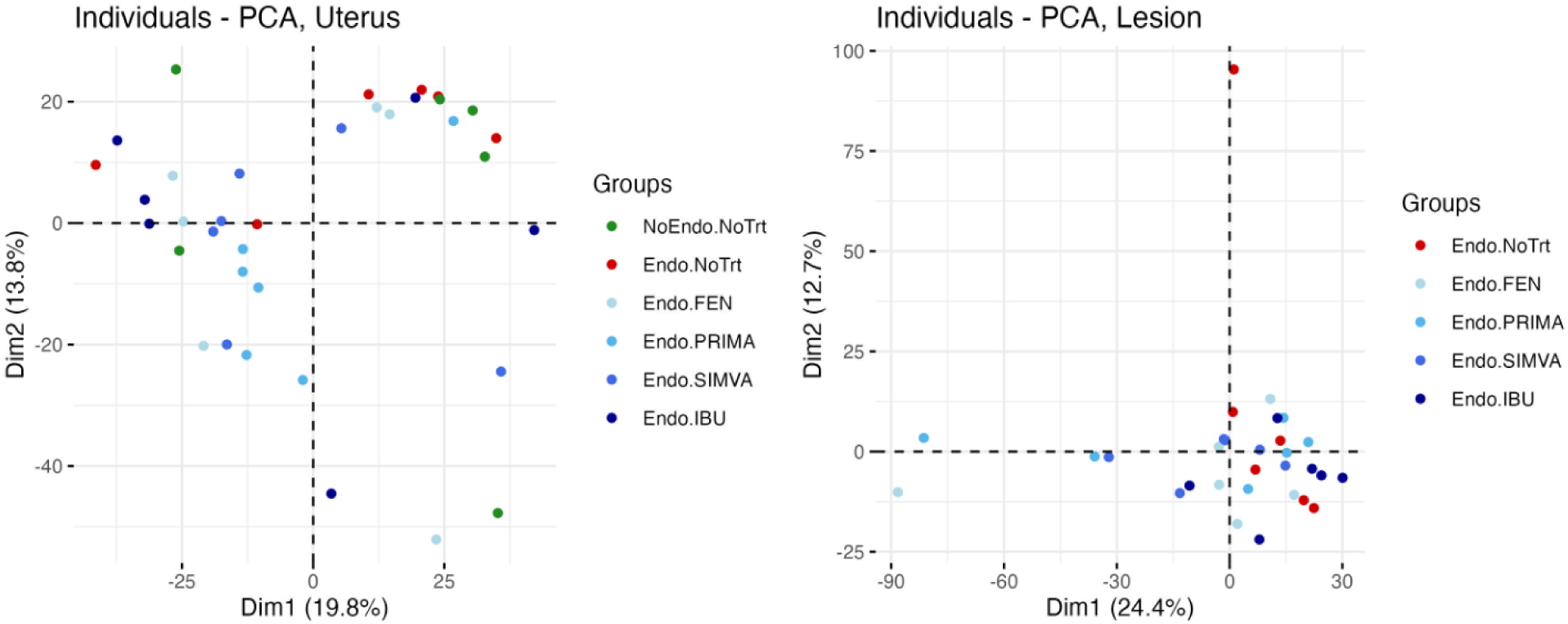
**Principal component analysis (PCA) for each sample source** -- uterus (left) and lesion (right). Each dot represents a sample.

**Fig. S3.**
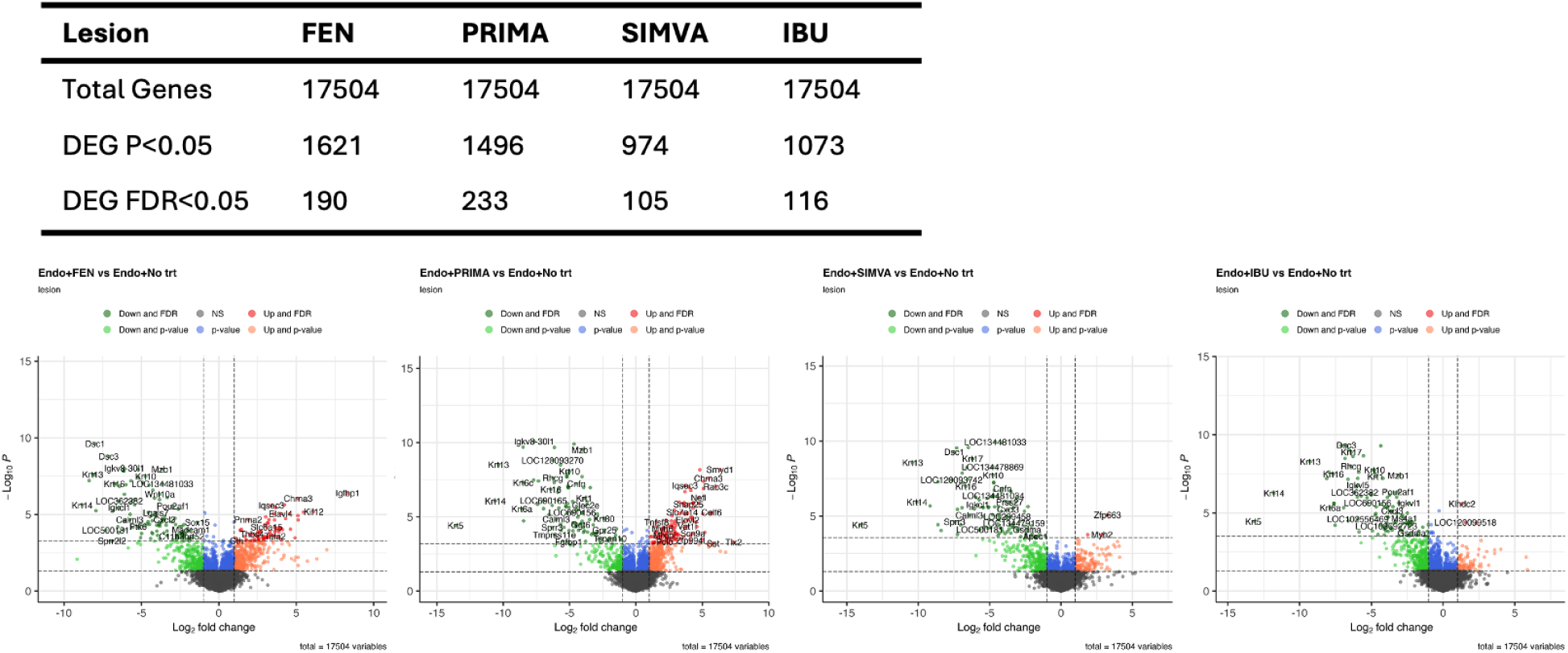
Volcano plots of individual differentially expressed genes (DEGs) in lesion samples by treatment group. The x-axis reflects the negative (-) log_10_ of the Benjamini-Hochberg (BH)-adjusted p-value, and the y-axis reflects the log_2_ fold change.

**Fig. S4.**
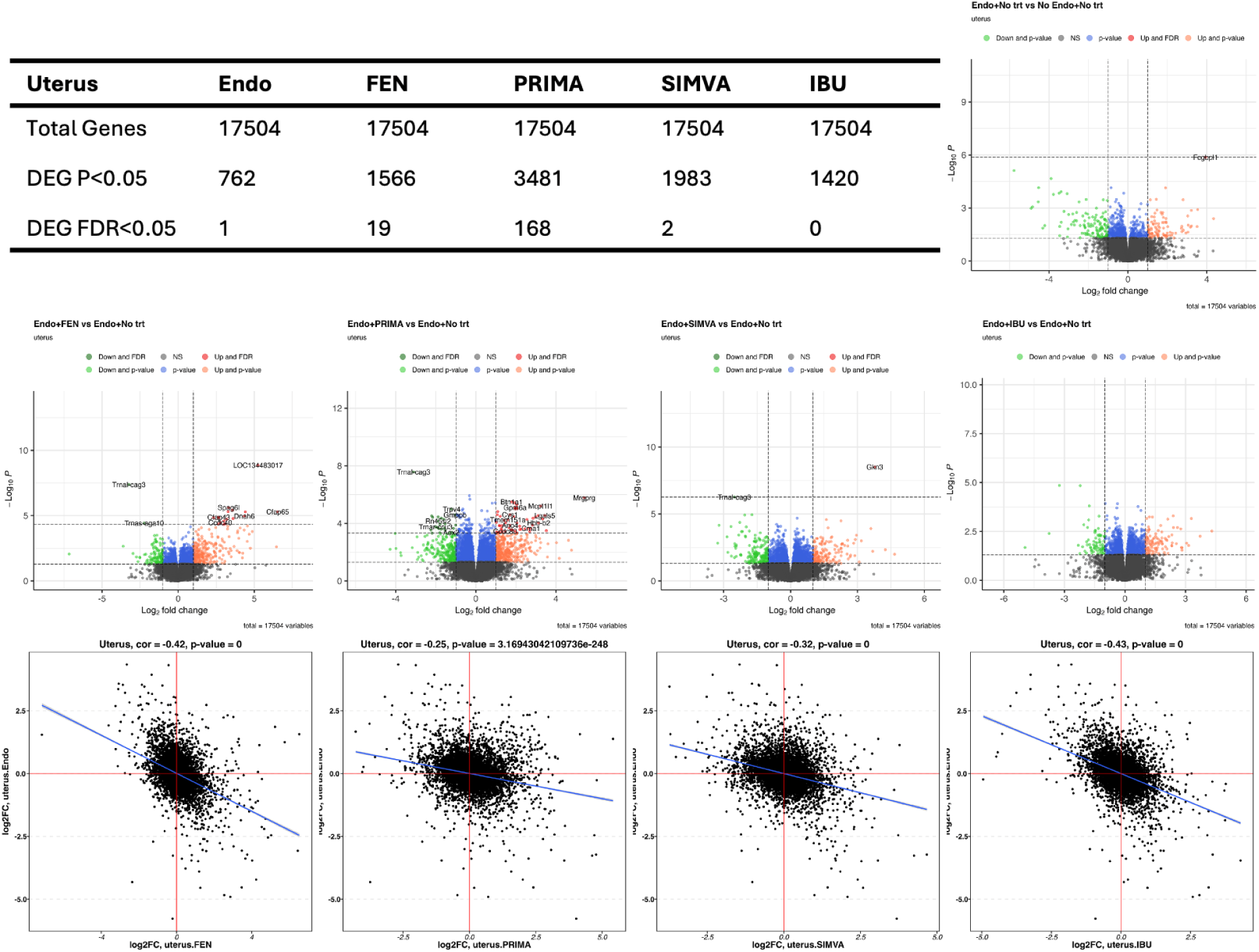
Volcano plots of individual differentially expressed genes (DEGs) in uterus samples by treatment group. The x-axis reflects the negative (-) log_10_ of the Benjamini-Hochberg (BH)-adjusted p-value, and the y-axis reflects the log_2_ fold change.

**Fig. S5.**
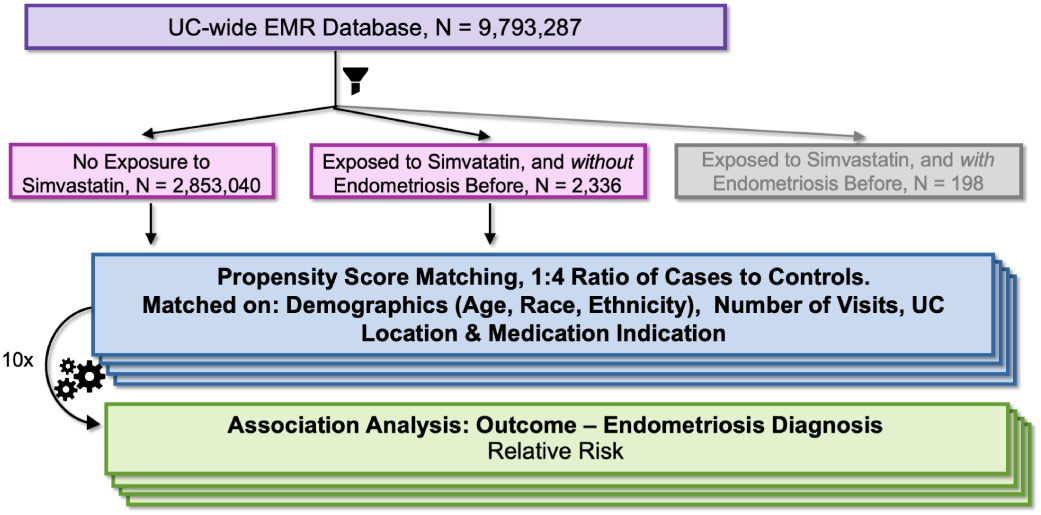
Overview of the Analysis for Simvastatin in Electronic Medical Records.

**Fig. S6.**
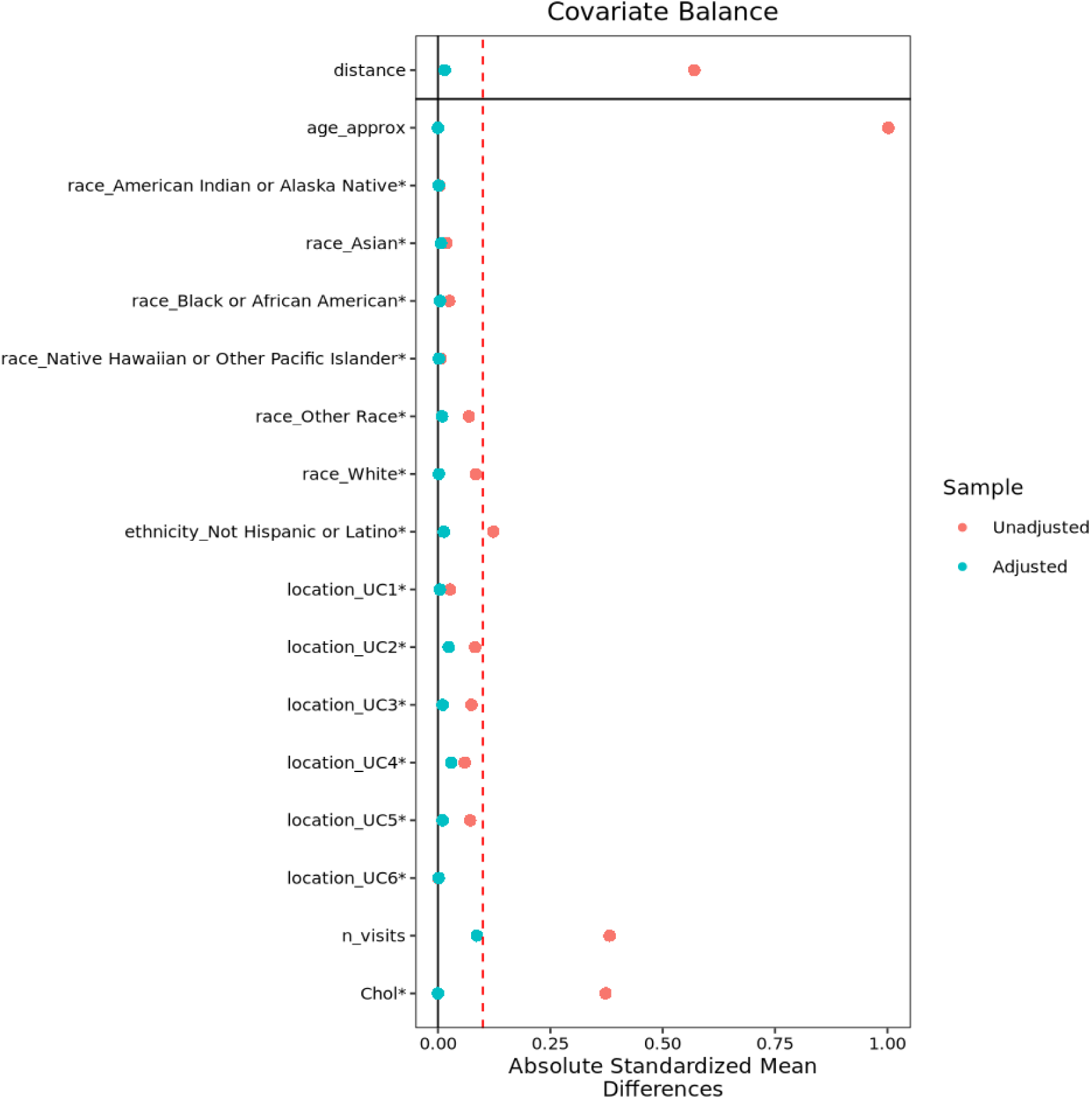
**Balance plot of the absolute standardized mean difference (SMD) of covariates before PS-matching (unadjusted, in pink) and after PS-matching (adjusted, in blue)**, with an absolute SMD of less than (0.1, reflected by the red-dashed line) indicative of adequate balance between groups.

## SUPPLEMENTARY TABLES

**Table S1.**
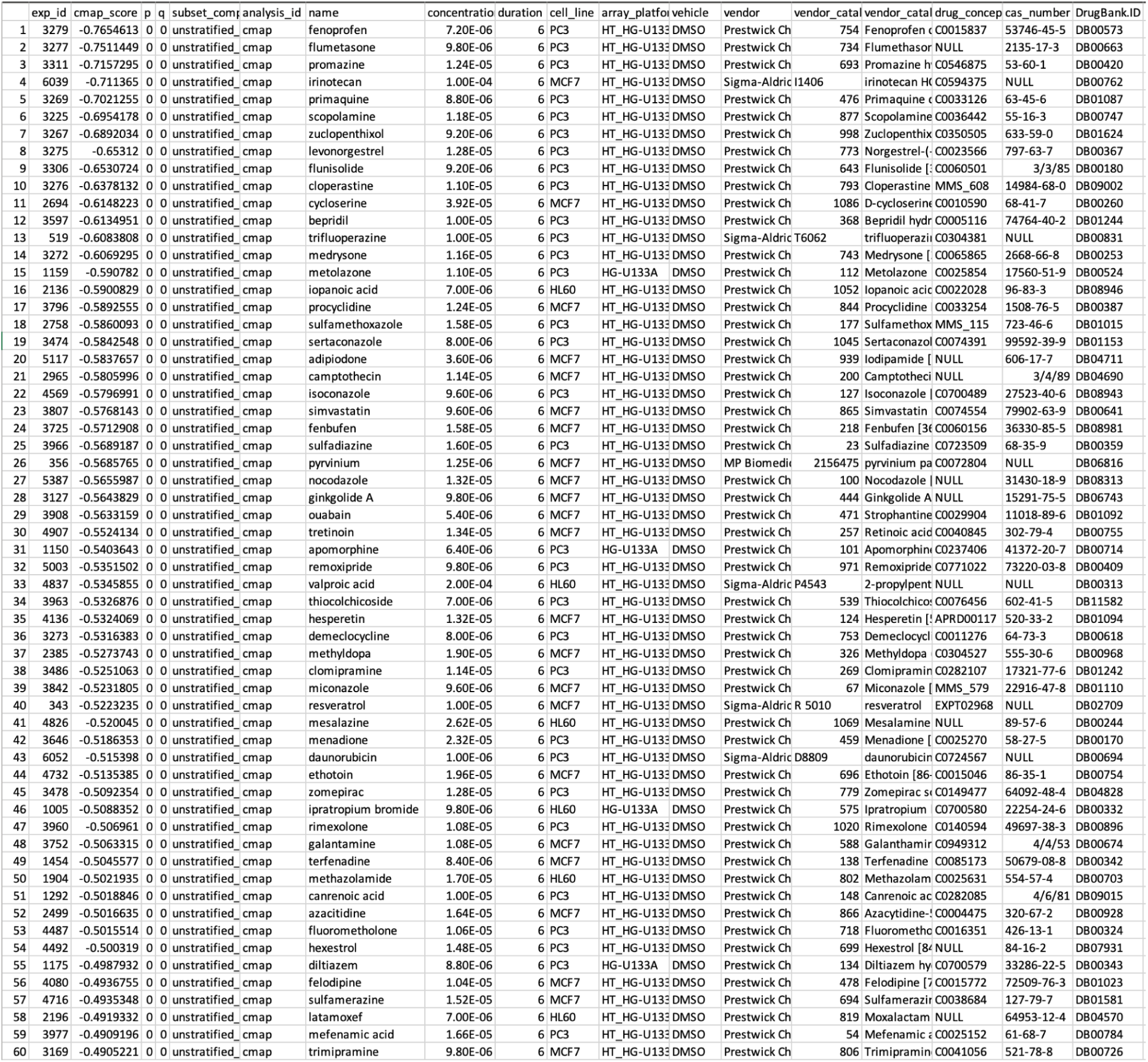
Drug candidates identified via computational drug repositioning pipeline using unstratified and stratified bulk transcriptomic signatures, https://ars.els-cdn.com/content/image/1-s2.0-S2589004224006096-mmc3.xlsx.

**Table S2. Differentially expressed genes (DEGs) in lesion samples in FEN treatment group.** lesion.FEN.csv

**Table S3. Differentially expressed genes (DEGs) in lesion samples in IBU treatment group.** lesion.IBU.csv

**Table S4. Differentially expressed genes (DEGs) in lesion samples in PRIMA treatment group.** lesion.PRIMA.csv

**Table S5. Differentially expressed genes (DEGs) in lesion samples in SIMVA treatment group.** lesion.SIMVA.csv

**Table S6. Differentially expressed genes (DEGs) in uterus samples in VEH treatment group.** uterus.Endo.csv

**Table S7. Differentially expressed genes (DEGs) in uterus samples in FEN treatment group.** uterus.FEN.csv

**Table S8. Differentially expressed genes (DEGs) in uterus samples in IBU treatment group.** uterus.IBU.csv

**Table S9. Differentially expressed genes (DEGs) in uterus samples in PRIMA treatment group.** uterus.PRIMA.csv

**Table S10. Differentially expressed genes (DEGs) in uterus samples in SIMVA treatment group.** uterus.SIMVA.csv

## Notes

### Summary of Updates

We have revised our manuscript to include results from our analysis of electronic medical records to validate simvastatin, one of the top drug repurposing candidates

