## supplementary_materials for "A Transcriptomics-Based Computational Drug Repurposing Pipeline Identifies Simvastatin And Primaquine As Therapeutics For Endometriosis"

### SUPPLEMENTARY FIGURES

| Volume (mL) | Baseline period (BL) median | BL IQR | Post-Endo period (PE) median | PE IQR | Post-Treatment period (PT) median | PT IQR | BL vs PE period MWU test Bonferroni-adjusted p-values | PE vs PT period MWU test Bonferroni-adjusted p-values | BL vs PT period MWU test Bonferroni-adjusted p-values |
|---|---|---|---|---|---|---|---|---|---|
| **CONTROL NO SURGERY (Negative control)** | | | | | | | | | |
| 0.01 | 0 | (0-0) | 0 | (0-0) | 0 | (0-0) | 1 | 1 | 1 |
| 0.15 | 0 | (0-0) | 0 | (0-0) | 0 | (0-0) | 1 | 1 | 1 |
| 0.30 | 0 | (0-0) | 0 | (0-33.3) | 0 | (0-33.3) | 1 | 1 | 1 |
| 0.40 | 33.3 | (0-33.3) | 33.3 | (0-33.3) | 33.3 | (0-33.3) | 1 | 1 | 1 |
| 0.55 | 33.3 | (33.3-33.3) | 33.3 | (33.3-66.6) | 33.3 | (33.3-66.6) | 1 | 1 | 1 |
| 0.70 | 66.6 | (33.3-66.6) | 66.6 | (66.6-66.6) | 66.6 | (66.6-66.6) | 1 | 1 | 1 |
| 0.80 | 100 | (66.6-100) | 100 | (66.6-100) | 100 | (66.6-100) | 1 | 1 | 1 |
| 0.90 | 100 | (100-100) | 100 | (100-100) | 100 | (100-100) | 1 | 1 | 1 |
| **VEHICLE (Negative control)** | | | | | | | | | |
| 0.01 | 0 | (0-0) | 0 | (0-0) | 0 | (0-0) | 1 | 1 | 1 |
| 0.15 | 0 | (0-0) | 33.3 | (33.3-66.6) | 33.3 | (33.3-33.3) | 1.7E-05 **** | 1 | 1.1E-05 **** |
| 0.30 | 0 | (0-33.3) | 66.6 | (33.3-66.6) | 66.6 | (33.3-66.6) | 9.8E-06 **** | 1 | 7.0E-06 **** |
| 0.40 | 33.3 | (0-33.3) | 66.6 | (66.6-100) | 66.6 | (66.6-66.6) | 4.1E-08 **** | 1 | 1.1E-06 **** |
| 0.55 | 33.3 | (33.3-41.6) | 100 | (66.6-100) | 100 | (75.0-100) | 1.7E-11 **** | 1 | 4.8E-09 **** |
| 0.70 | 66.6 | (58.3-75.0) | 100 | (100-100) | 100 | (100-100) | 4.3E-06 **** | 1 | 2.3E-05 **** |
| 0.80 | 100 | (66.6-100) | 100 | (100-100) | 100 | (100-100) | 6.1E-05 **** | 1 | 0.02 * |
| 0.90 | 100 | (100-100) | 100 | (100-100) | 100 | (100-100) | 1 | 1 | 1 |
| **IBUPROFEN (Positive control)** | | | | | | | | | |
| 0.01 | 0 | (0-0) | 0 | (0-0) | 0 | (0-0) | 1 | 1 | 1 |
| 0.15 | 0 | (0-0) | 16.65 | (0-33.3) | 0 | (0-0) | 3.5E-03 ** | 2.3E-03 ** | 1 |
| 0.30 | 0 | (0-0) | 33.3 | (0-66.6) | 0 | (0-0) | 2.0E-03 ** | 9.7E-04 ** | 1 |
| 0.40 | 0 | (0-25) | 66.6 | (33.3-66.6) | 0 | (0-33.3) | 1.5E-05 **** | 2.9E-06 **** | 1 |
| 0.55 | 33.3 | (8.3-33.3) | 83.3 | (66.6-100) | 0 | (0-33.3) | 2.5E-07 **** | 3.4E-08 **** | 1 |
| 0.70 | 66.6 | (66.6-66.6) | 100 | (100-100) | 66.6 | (33.3-66.6) | 5.8E-05 **** | 5.1E-06 **** | 1 |
| 0.80 | 100 | (66.6-100) | 100 | (100-100) | 100 | (66.6-100) | 3.9E-03 ** | 1.8E-03 ** | 1 |
| 0.90 | 100 | (74.9-100) | 100 | (100-100) | 100 | (100-100) | 0.12 | 0.8 | 1 |
| **FENOPROFEN** | | | | | | | | | |
| 0.01 | 0 | (0-0) | 0 | (0-0) | 0 | (0-0) | 1 | 1 | 1 |
| 0.15 | 0 | (0-0) | 33.3 | (0-66.6) | 0 | (0-0) | 2.7E-04 *** | 2.3E-03 ** | 0.61 |
| 0.30 | 0 | (0-25) | 33.3 | (33.3-66.6) | 0 | (0-33.3) | 2.0E-05 **** | 3.0E-06 **** | 1 |
| 0.40 | 0 | (0-33.3) | 66.6 | (33.3-91.7) | 16.65 | (0-33.3) | 4.0E-06 **** | 1.8E-05 **** | 1 |
| 0.55 | 33.3 | (8.3-33.3) | 83.3 | (66.6-100) | 33.3 | (33.3-66.6) | 2.7E-07 **** | 7.9E-07 **** | 0.16 |
| 0.70 | 66.6 | (33.3-66.6) | 100 | (100-100) | 66.6 | (66.6-100) | 1.5E-03 ** | 4.4E-02 * | 0.97 |
| 0.80 | 66.6 | (66.6-100) | 100 | (100-100) | 100 | (66.6-100) | 4.8E-05 **** | 5.8E-03 * | 1 |
| 0.90 | 100 | (100-100) | 100 | (100-100) | 100 | (100-100) | 0.34 | 0.8 | 1 |
| **PRIMAQUINE** | | | | | | | | | |
| 0.01 | 0 | (0-0) | | (0-0) | 0 | (0-0) | 1 | 1 | 1 |
| 0.15 | 0 | (0-0) | 33.3 | (0-66.6) | 0 | (0-33.3) | 3.7E-06 **** | 1.4E-03 ** | 0.24 |
| 0.30 | 0 | (0-0) | 50 | (33.3-66.6) | 33.3 | (0-33.3) | 1.2E-07 **** | 1.7E-02 * | 2.6E-03 ** |
| 0.40 | 33.3 | (0-33.3) | 66.6 | (41.6-100) | 50 | (33.3-66.6) | 4.5E-09 **** | 3.3E-02 * | 1.6E-05 **** |
| 0.55 | 33.3 | (33.3-41.6) | 100 | (66.6-100) | 66.6 | (33.3-100) | 3.4E-10 **** | 1.0E-03 ** | 0.03 * |
| 0.70 | 50 | (33.3-66.6) | 100 | (100-100) | 66.6 | (66.6-100) | 6.5E-09 **** | 5.8E-03 * | 9.8E-04 ** |
| 0.80 | 66.6 | (66.6-100) | 100 | (100-100) | 100 | (66.6-100) | 6.2E-03 * | 9.3E-02 | 1 |
| 0.90 | 100 | (100-100) | 100 | (100-100) | 100 | (100-100) | 0.47 | 1 | 1 |
| **SIMVASTATIN** | | | | | | | | | |
| 0.01 | 0 | (0-0) | 0 | (0-0) | 0 | (0-0) | 1 | 1 | 1 |
| 0.15 | 0 | (0-0) | 33.3 | (33.3-66.6) | 0 | (0-0) | 6.8E-08 **** | 1.7E-09 **** | 1 |
| 0.30 | 0 | (0-8.3) | 66.6 | (41.6-66.6) | 0 | (0-25.0) | 2.0E-11 **** | 1.0E-12 **** | 1 |
| 0.40 | 33.3 | (0-33.3) | 66.6 | (66.6-100) | 33.3 | (33.3-33.3) | 6.0E-11 **** | 3.9E-08 **** | 0.08 |
| 0.55 | 33.3 | (33.3-66.6) | 100 | (66.6-100) | 33.3 | (33.3-66.6) | 5.4E-09 **** | 1.1E-07 **** | 1 |
| 0.70 | 66.6 | (33.3-66.6) | 100 | (100-100) | 66.6 | (41.6-100) | 5.0E-07 **** | 2.3E-04 *** | 1 |
| 0.80 | 83.3 | (66.6-100) | 100 | (100-100) | 83.3 | (66.6-100) | 9.0E-05 **** | 1.5E-05 **** | 1 |
| 0.90 | 100 | (100-100) | 100 | (100-100) | 100 | (100-100) | 1 | 1 | 1 |

**Fig. S1: Endometriosis animal model escape responses.** Median escape response with interquartile range (IQR) for each delivered volume (0.01, 0.15,0.30, 0.40, 0.55, 0.70, 0.80, and 0.90 mL) during the baseline, post-endo surgery, and post-treatment periods, with Bonferroni-corrected p-values from Mann-Whitney U test for baseline period vs post-endo surgery period, post-endo surgery vs post-treatment period, and baseline period vs post-treatment period (* denotes <0.05, ** denotes <0.005, *** denotes <0.0005, and **** denotes <0.0001). Endo rats (n=6/group) vaginal nociception assessed. Treated with vehicle (negative control), primaquine, simvastatin, fenoprofen, or ibuprofen (positive control). Note: Fenoprofen and ibuprofen data are from prior work*(23)*

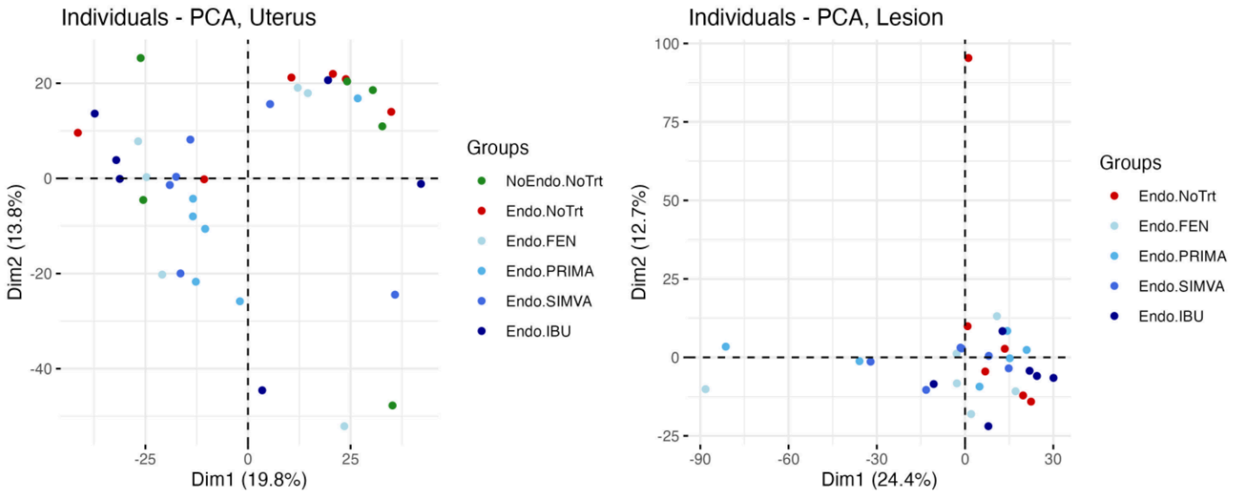


**Fig. S2. Principal component analysis (PCA) for each sample source** -- uterus (left) and lesion (right). Each dot represents a sample.

| Lesion | FEN | PRIMA | SIMVA | IBU |
|---|---|---|---|---|
| Total Genes | 17504 | 17504 | 17504 | 17504 |
| DEG P<0.05 | 1621 | 1496 | 974 | 1073 |
| DEG FDR<0.05 | 190 | 233 | 105 | 116 |

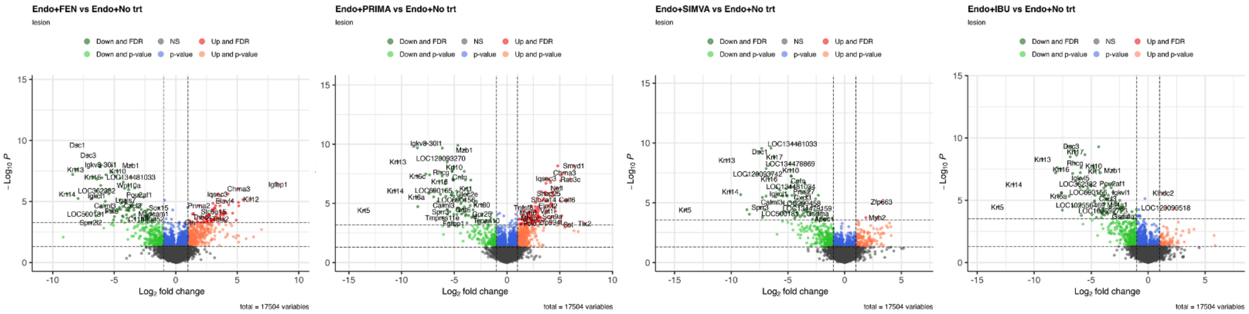


**Fig. S3. Volcano plots of individual differentially expressed genes (DEGs) in lesion samples by treatment group**. The x-axis reflects the negative (-) $\log_{10}$ of the Benjamini-Hochberg (BH)-adjusted p-value, and the y-axis reflects the $\log_2$ fold change.

| Uterus | Endo | FEN | PRIMA | SIMVA | IBU |
|---|---|---|---|---|---|
| Total Genes | 17504 | 17504 | 17504 | 17504 | 17504 |
| DEG P<0.05 | 762 | 1566 | 3481 | 1983 | 1420 |
| DEG FDR<0.05 | 1 | 19 | 168 | 2 | 0 |

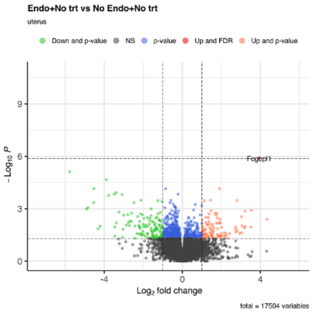


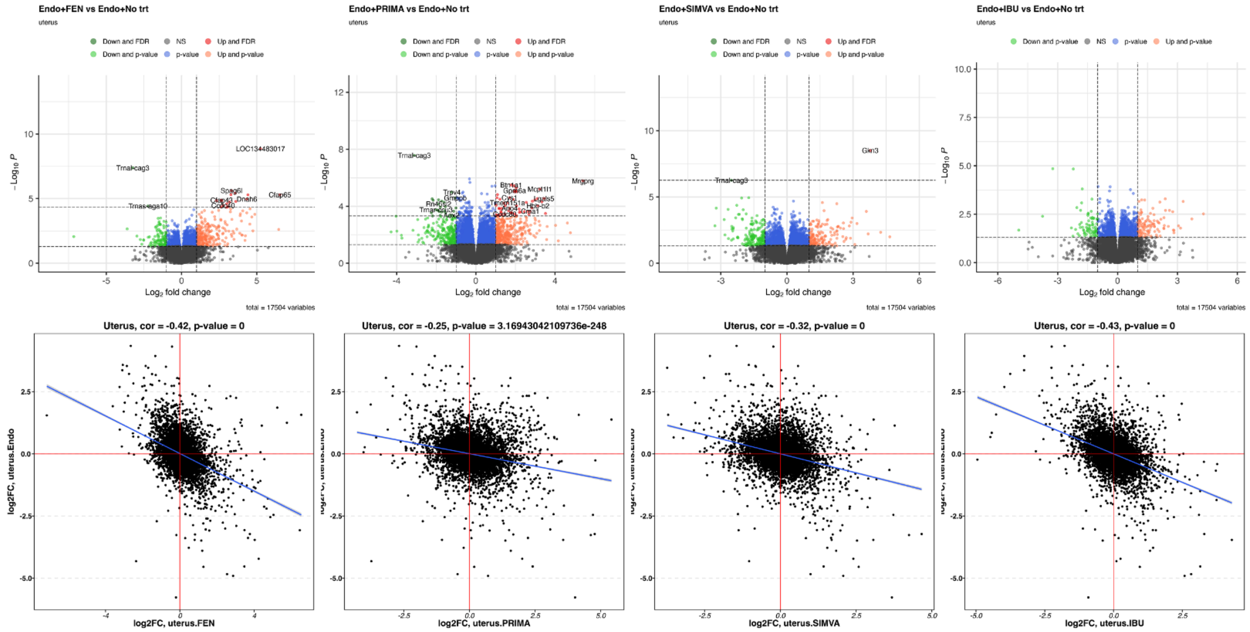


**Fig. S4. Volcano plots of individual differentially expressed genes (DEGs) in uterus samples by treatment group.** The x-axis reflects the negative (-) $\log_{10}$ of the Benjamini-Hochberg (BH)-adjusted p-value, and the y-axis reflects the $\log_2$ fold change.

**Fig. S5. Overview of the Analysis for Simvastatin in Electronic Medical Records**

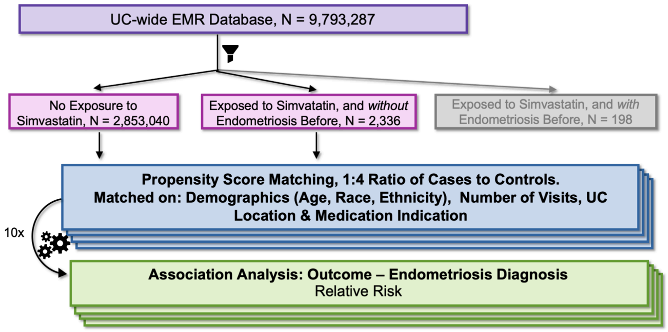


**Fig. S6. Balance plot of the absolute standardized mean difference (SMD) of covariates before PS-matching (unadjusted, in pink) and after PS-matching (adjusted, in blue)**, with an absolute SMD of less than (0.1, reflected by the red-dashed line) indicative of adequate balance between groups.

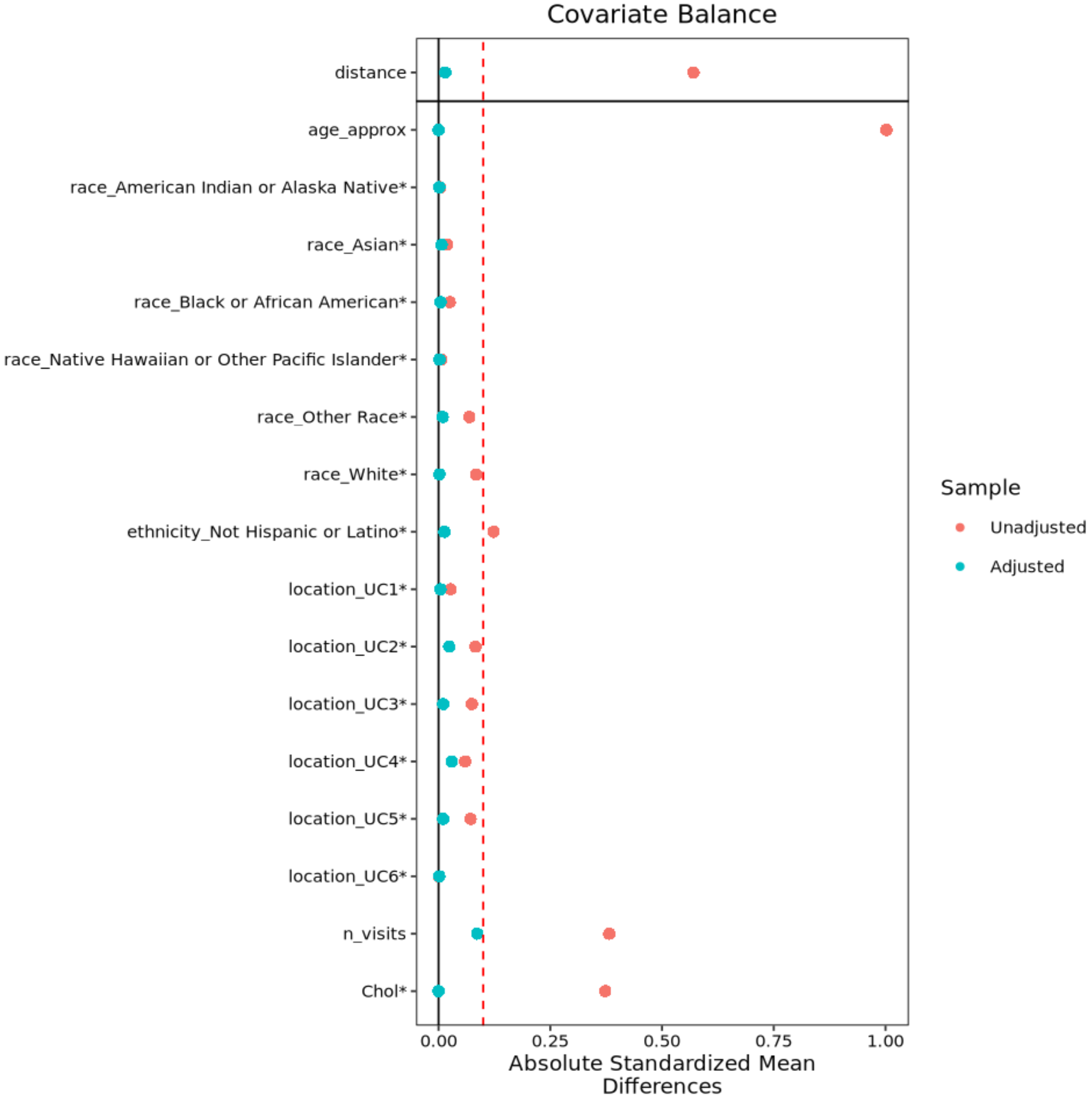

### SUPPLEMENTARY TABLES

**Table S1. Drug candidates identified via computational drug repositioning pipeline using unstratified and stratified bulk transcriptomic signatures, https://ars.els-cdn.com/content/image/1-s2.0-S2589004224006096-mmc3.xlsx**

| | exp_id | cmap_score | p | q | subset_comp | analysis_id | name | concentratio | duration | cell_line | array_platfo | vehicle | vendor | vendor_catal | vendor_catal | drug_concep | cas_number | DrugBank.ID |
|---|---|---|---|---|---|---|---|---|---|---|---|---|---|---|---|---|---|---|
| 1 | 3279 | -0.7654613 | 0 | 0 | unstratified_ | cmap | fenoprofen | 7.20E-06 | 6 | PC3 | HT_HG-U133 | DMSO | Prestwick Ch | 754 | Fenoprofen ( | C0015837 | 53746-45-5 | DB00573 |
| 2 | 3277 | -0.7511449 | 0 | 0 | unstratified_ | cmap | flumetasone | 9.80E-06 | 6 | PC3 | HT_HG-U133 | DMSO | Prestwick Ch | 734 | Flumethasor | NULL | 2135-17-3 | DB00663 |
| 3 | 3311 | -0.7157295 | 0 | 0 | unstratified_ | cmap | promazine | 1.24E-05 | 6 | PC3 | HT_HG-U133 | DMSO | Prestwick Ch | 693 | Promazine h | C0546875 | 53-60-1 | DB00420 |
| 4 | 6039 | -0.711365 | 0 | 0 | unstratified_ | cmap | irinotecan | 1.00E-04 | 6 | MCF7 | HT_HG-U133 | DMSO | Sigma-Aldric | I1406 | irinotecan H | C0594375 | NULL | DB00762 |
| 5 | 3269 | -0.7021255 | 0 | 0 | unstratified_ | cmap | primaquine | 8.80E-06 | 6 | PC3 | HT_HG-U133 | DMSO | Prestwick Ch | 476 | Primaquine ( | C0033126 | 63-45-6 | DB01087 |
| 6 | 3225 | -0.6954178 | 0 | 0 | unstratified_ | cmap | scopolamine | 1.18E-05 | 6 | PC3 | HT_HG-U133 | DMSO | Prestwick Ch | 877 | Scopolamine | C0036442 | 55-16-3 | DB00747 |
| 7 | 3267 | -0.6892034 | 0 | 0 | unstratified_ | cmap | zuclopenthixol | 9.20E-06 | 6 | PC3 | HT_HG-U133 | DMSO | Prestwick Ch | 998 | Zuclopenthix | C0350505 | 633-59-0 | DB01624 |
| 8 | 3275 | -0.65312 | 0 | 0 | unstratified_ | cmap | levonorgestrel | 1.28E-05 | 6 | PC3 | HT_HG-U133 | DMSO | Prestwick Ch | 773 | Norgestrel-(- | C0023566 | 797-63-7 | DB00367 |
| 9 | 3306 | -0.6530724 | 0 | 0 | unstratified_ | cmap | flunisolide | 9.20E-06 | 6 | PC3 | HT_HG-U133 | DMSO | Prestwick Ch | 643 | Flunisolide [ | C0060501 | 3/3/85 | DB00180 |
| 10 | 3276 | -0.6378132 | 0 | 0 | unstratified_ | cmap | cloperastine | 1.10E-05 | 6 | PC3 | HT_HG-U133 | DMSO | Prestwick Ch | 793 | Cloperastine | MMS_608 | 14984-68-0 | DB09002 |
| 11 | 2694 | -0.6148223 | 0 | 0 | unstratified_ | cmap | cycloserine | 3.92E-05 | 6 | MCF7 | HT_HG-U133 | DMSO | Prestwick Ch | 1086 | D-cycloserine | C0010590 | 68-41-7 | DB00260 |
| 12 | 3597 | -0.6134951 | 0 | 0 | unstratified_ | cmap | bepridil | 1.00E-05 | 6 | PC3 | HT_HG-U133 | DMSO | Prestwick Ch | 368 | Bepridil hydr | C0005116 | 74764-40-2 | DB01244 |
| 13 | 519 | -0.6083808 | 0 | 0 | unstratified_ | cmap | trifluoperazine | 1.00E-05 | 6 | PC3 | HT_HG-U133 | DMSO | Sigma-Aldric | T6062 | trifluoperazi | C0304381 | NULL | DB00831 |
| 14 | 3272 | -0.6069295 | 0 | 0 | unstratified_ | cmap | medrysone | 1.16E-05 | 6 | PC3 | HT_HG-U133 | DMSO | Prestwick Ch | 743 | Medrysone [ | C0065865 | 2668-66-8 | DB00253 |
| 15 | 1159 | -0.590782 | 0 | 0 | unstratified_ | cmap | metolazone | 1.10E-05 | 6 | PC3 | HG-U133A | DMSO | Prestwick Ch | 112 | Metolazone | C0025854 | 17560-51-9 | DB00524 |
| 16 | 2136 | -0.5900829 | 0 | 0 | unstratified_ | cmap | iopanoic acid | 7.00E-06 | 6 | HL60 | HT_HG-U133 | DMSO | Prestwick Ch | 1052 | Iopanoic acic | C0022028 | 96-83-3 | DB08946 |
| 17 | 3796 | -0.5892555 | 0 | 0 | unstratified_ | cmap | procyclidine | 1.24E-05 | 6 | MCF7 | HT_HG-U133 | DMSO | Prestwick Ch | 844 | Procyclidine | C0033254 | 1508-76-5 | DB00387 |
| 18 | 2758 | -0.5860093 | 0 | 0 | unstratified_ | cmap | sulfamethoxazole | 1.58E-05 | 6 | PC3 | HT_HG-U133 | DMSO | Prestwick Ch | 177 | Sulfamethox | MMS_115 | 723-46-6 | DB01015 |
| 19 | 3474 | -0.5842548 | 0 | 0 | unstratified_ | cmap | sertaconazole | 8.00E-06 | 6 | PC3 | HT_HG-U133 | DMSO | Prestwick Ch | 1045 | Sertaconazol | C0074391 | 99592-39-9 | DB01153 |
| 20 | 5117 | -0.5837657 | 0 | 0 | unstratified_ | cmap | adipiodone | 3.60E-06 | 6 | MCF7 | HT_HG-U133 | DMSO | Prestwick Ch | 939 | Iodipamide [ | NULL | 606-17-7 | DB04711 |
| 21 | 2965 | -0.5805996 | 0 | 0 | unstratified_ | cmap | camptothecin | 1.14E-05 | 6 | MCF7 | HT_HG-U133 | DMSO | Prestwick Ch | 200 | Camptotheci | NULL | 3/4/89 | DB04690 |
| 22 | 4569 | -0.5796991 | 0 | 0 | unstratified_ | cmap | isoconazole | 9.60E-06 | 6 | PC3 | HT_HG-U133 | DMSO | Prestwick Ch | 127 | Isoconazole | | C0700489 | 27523-40-6 | DB08943 |
| 23 | 3807 | -0.5768143 | 0 | 0 | unstratified_ | cmap | simvastatin | 9.60E-06 | 6 | MCF7 | HT_HG-U133 | DMSO | Prestwick Ch | 865 | Simvastatin | C0074554 | 79902-63-9 | DB00641 |
| 24 | 3725 | -0.5712908 | 0 | 0 | unstratified_ | cmap | fenbufen | 1.58E-05 | 6 | MCF7 | HT_HG-U133 | DMSO | Prestwick Ch | 218 | Fenbufen [36 | C0060156 | 36330-85-5 | DB08981 |
| 25 | 3966 | -0.5689187 | 0 | 0 | unstratified_ | cmap | sulfadiazine | 1.60E-05 | 6 | PC3 | HT_HG-U133 | DMSO | Prestwick Ch | 23 | Sulfadiazine | C0723509 | 68-35-9 | DB00359 |
| 26 | 356 | -0.5685765 | 0 | 0 | unstratified_ | cmap | pyrvinium | 1.25E-06 | 6 | MCF7 | HT_HG-U133 | DMSO | MP Biomedi | 2156475 | pyrvinium pa | C0072804 | NULL | DB06816 |
| 27 | 5387 | -0.5655987 | 0 | 0 | unstratified_ | cmap | nocodazole | 1.32E-05 | 6 | MCF7 | HT_HG-U133 | DMSO | Prestwick Ch | 100 | Nocodazole | | NULL | 31430-18-9 | DB08313 |
| 28 | 3127 | -0.5643829 | 0 | 0 | unstratified_ | cmap | ginkgolide A | 9.80E-06 | 6 | MCF7 | HT_HG-U133 | DMSO | Prestwick Ch | 444 | Ginkgolide A | NULL | 15291-75-5 | DB06743 |
| 29 | 3908 | -0.5633159 | 0 | 0 | unstratified_ | cmap | ouabain | 5.40E-06 | 6 | MCF7 | HT_HG-U133 | DMSO | Prestwick Ch | 471 | Strophantine | C0029904 | 11018-89-6 | DB01092 |
| 30 | 4907 | -0.5524134 | 0 | 0 | unstratified_ | cmap | tretinoin | 1.34E-05 | 6 | MCF7 | HT_HG-U133 | DMSO | Prestwick Ch | 257 | Retinoic acid | C0040845 | 302-79-4 | DB00755 |
| 31 | 1150 | -0.5403643 | 0 | 0 | unstratified_ | cmap | apomorphine | 6.40E-06 | 6 | PC3 | HG-U133A | DMSO | Prestwick Ch | 101 | Apomorphine | C0237406 | 41372-20-7 | DB00714 |
| 32 | 5003 | -0.5351502 | 0 | 0 | unstratified_ | cmap | remoxipride | 9.80E-06 | 6 | PC3 | HT_HG-U133 | DMSO | Prestwick Ch | 971 | Remoxipride | C0771022 | 73220-03-8 | DB00409 |
| 33 | 4837 | -0.5345855 | 0 | 0 | unstratified_ | cmap | valproic acid | 2.00E-04 | 6 | HL60 | HT_HG-U133 | DMSO | Sigma-Aldric | P4543 | 2-propylpent | NULL | NULL | DB00313 |
| 34 | 3963 | -0.5326876 | 0 | 0 | unstratified_ | cmap | thiocolchicoside | 7.00E-06 | 6 | PC3 | HT_HG-U133 | DMSO | Prestwick Ch | 539 | Thiocolchicos | C0076456 | 602-41-5 | DB11582 |
| 35 | 4136 | -0.5324069 | 0 | 0 | unstratified_ | cmap | hesperetin | 1.32E-05 | 6 | MCF7 | HT_HG-U133 | DMSO | Prestwick Ch | 124 | Hesperetin [ | APRD00117 | 520-33-2 | DB01094 |
| 36 | 3273 | -0.5316383 | 0 | 0 | unstratified_ | cmap | demeclocycline | 8.00E-06 | 6 | PC3 | HT_HG-U133 | DMSO | Prestwick Ch | 753 | Demeclocycl | C0011276 | 64-73-3 | DB00618 |
| 37 | 2385 | -0.5273743 | 0 | 0 | unstratified_ | cmap | methyldopa | 1.90E-05 | 6 | MCF7 | HT_HG-U133 | DMSO | Prestwick Ch | 326 | Methyldopa | C0304527 | 555-30-6 | DB00968 |
| 38 | 3486 | -0.5251063 | 0 | 0 | unstratified_ | cmap | clomipramine | 1.14E-05 | 6 | PC3 | HT_HG-U133 | DMSO | Prestwick Ch | 269 | Clomipramin | C0282107 | 17321-77-6 | DB01242 |
| 39 | 3842 | -0.5231805 | 0 | 0 | unstratified_ | cmap | miconazole | 9.60E-06 | 6 | MCF7 | HT_HG-U133 | DMSO | Prestwick Ch | 67 | Miconazole [ | MMS_579 | 22916-47-8 | DB01110 |
| 40 | 343 | -0.5223235 | 0 | 0 | unstratified_ | cmap | resveratrol | 1.00E-05 | 6 | MCF7 | HT_HG-U133 | DMSO | Sigma-Aldric | R 5010 | resveratrol | EXPT02968 | NULL | DB02709 |
| 41 | 4826 | -0.520045 | 0 | 0 | unstratified_ | cmap | mesalazine | 2.62E-05 | 6 | HL60 | HT_HG-U133 | DMSO | Prestwick Ch | 1069 | Mesalamine | NULL | 89-57-6 | DB00244 |
| 42 | 3646 | -0.5186353 | 0 | 0 | unstratified_ | cmap | menadione | 2.32E-05 | 6 | PC3 | HT_HG-U133 | DMSO | Prestwick Ch | 459 | Menadione [ | C0025270 | 58-27-5 | DB00170 |
| 43 | 6052 | -0.515398 | 0 | 0 | unstratified_ | cmap | daunorubicin | 1.00E-06 | 6 | PC3 | HT_HG-U133 | DMSO | Sigma-Aldric | D8809 | daunorubicin | C0724567 | NULL | DB00694 |
| 44 | 4732 | -0.5135385 | 0 | 0 | unstratified_ | cmap | ethotoin | 1.96E-05 | 6 | MCF7 | HT_HG-U133 | DMSO | Prestwick Ch | 696 | Ethotoin [86- | C0015046 | 86-35-1 | DB00754 |
| 45 | 3478 | -0.5092354 | 0 | 0 | unstratified_ | cmap | zomepirac | 1.28E-05 | 6 | PC3 | HT_HG-U133 | DMSO | Prestwick Ch | 779 | Zomepirac s | C0149477 | 64092-48-4 | DB04828 |
| 46 | 1005 | -0.5088352 | 0 | 0 | unstratified_ | cmap | ipratropium bromide | 9.80E-06 | 6 | HL60 | HG-U133A | DMSO | Prestwick Ch | 575 | Ipratropium | C0700580 | 22254-24-6 | DB00332 |
| 47 | 3960 | -0.506961 | 0 | 0 | unstratified_ | cmap | rimexolone | 1.08E-05 | 6 | PC3 | HT_HG-U133 | DMSO | Prestwick Ch | 1020 | Rimexolone | C0140594 | 49697-38-3 | DB00896 |
| 48 | 3752 | -0.5063315 | 0 | 0 | unstratified_ | cmap | galantamine | 1.08E-05 | 6 | MCF7 | HT_HG-U133 | DMSO | Prestwick Ch | 588 | Galanthamir | C0949312 | 4/4/53 | DB00674 |
| 49 | 1454 | -0.5045577 | 0 | 0 | unstratified_ | cmap | terfenadine | 8.40E-06 | 6 | MCF7 | HT_HG-U133 | DMSO | Prestwick Ch | 138 | Terfenadine | C0085173 | 50679-08-8 | DB00342 |
| 50 | 1904 | -0.5021935 | 0 | 0 | unstratified_ | cmap | methazolamide | 1.70E-05 | 6 | HL60 | HT_HG-U133 | DMSO | Prestwick Ch | 802 | Methazolam | C0025631 | 554-57-4 | DB00703 |
| 51 | 1292 | -0.5018846 | 0 | 0 | unstratified_ | cmap | canrenoic acid | 1.00E-05 | 6 | PC3 | HT_HG-U133 | DMSO | Prestwick Ch | 148 | Canrenoic ac | C0282085 | 4/6/81 | DB09015 |
| 52 | 2499 | -0.5016635 | 0 | 0 | unstratified_ | cmap | azacitidine | 1.64E-05 | 6 | MCF7 | HT_HG-U133 | DMSO | Prestwick Ch | 866 | Azacytidine- | C0004475 | 320-67-2 | DB00928 |
| 53 | 4487 | -0.5015514 | 0 | 0 | unstratified_ | cmap | fluorometholone | 1.06E-05 | 6 | PC3 | HT_HG-U133 | DMSO | Prestwick Ch | 718 | Fluorometho | C0016351 | 426-13-1 | DB00324 |
| 54 | 4492 | -0.500319 | 0 | 0 | unstratified_ | cmap | hexestrol | 1.48E-05 | 6 | PC3 | HT_HG-U133 | DMSO | Prestwick Ch | 699 | Hexestrol [84 | NULL | 84-16-2 | DB07931 |
| 55 | 1175 | -0.4987932 | 0 | 0 | unstratified_ | cmap | diltiazem | 8.80E-06 | 6 | PC3 | HG-U133A | DMSO | Prestwick Ch | 134 | Diltiazem hy | C0700579 | 33286-22-5 | DB00343 |
| 56 | 4080 | -0.4936755 | 0 | 0 | unstratified_ | cmap | felodipine | 1.04E-05 | 6 | MCF7 | HT_HG-U133 | DMSO | Prestwick Ch | 478 | Felodipine [7 | C0015772 | 72509-76-3 | DB01023 |
| 57 | 4716 | -0.4935348 | 0 | 0 | unstratified_ | cmap | sulfamerazine | 1.52E-05 | 6 | MCF7 | HT_HG-U133 | DMSO | Prestwick Ch | 694 | Sulfamerazir | C0038684 | 127-79-7 | DB01581 |
| 58 | 2196 | -0.4919332 | 0 | 0 | unstratified_ | cmap | latamoxef | 7.00E-06 | 6 | HL60 | HT_HG-U133 | DMSO | Prestwick Ch | 819 | Moxalactam | NULL | 64953-12-4 | DB04570 |
| 59 | 3977 | -0.4909196 | 0 | 0 | unstratified_ | cmap | mefenamic acid | 1.66E-05 | 6 | PC3 | HT_HG-U133 | DMSO | Prestwick Ch | 54 | Mefenamic a | C0025152 | 61-68-7 | DB00784 |
| 60 | 3169 | -0.4905221 | 0 | 0 | unstratified_ | cmap | trimipramine | 9.80E-06 | 6 | MCF7 | HT_HG-U133 | DMSO | Prestwick Ch | 806 | Trimipramin | C0041056 | 521-78-8 | DB00726 |

**Table S2. Differentially expressed genes (DEGs) in lesion samples in FEN treatment group.** lesion.FEN.csv

**Table S3. Differentially expressed genes (DEGs) in lesion samples in IBU treatment group.** lesion.IBU.csv

**Table S4. Differentially expressed genes (DEGs) in lesion samples in PRIMA treatment group.** lesion.PRIMA.csv

**Table S5. Differentially expressed genes (DEGs) in lesion samples in SIMVA treatment group.** lesion.SIMVA.csv

**Table S6. Differentially expressed genes (DEGs) in uterus samples in VEH treatment group.** uterus.Endo.csv

**Table S7. Differentially expressed genes (DEGs) in uterus samples in FEN treatment group.** uterus.FEN.csv

**Table S8. Differentially expressed genes (DEGs) in uterus samples in IBU treatment group.** uterus.IBU.csv

**Table S9. Differentially expressed genes (DEGs) in uterus samples in PRIMA treatment group.** uterus.PRIMA.csv

**Table S10. Differentially expressed genes (DEGs) in uterus samples in SIMVA treatment group.** uterus.SIMVA.csv
